# A novel flight style allowing the smallest featherwing beetles to excel

**DOI:** 10.1101/2021.06.24.449497

**Authors:** Sergey E. Farisenkov, Dmitry Kolomenskiy, Pyotr N. Petrov, Nadejda A. Lapina, Thomas Engels, Fritz-Olaf Lehmann, Ryo Onishi, Hao Liu, Alexey A. Polilov

## Abstract

Flight speed generally correlates positively with animal body size^1^. Surprisingly, miniature featherwing beetles can fly at speeds and accelerations of insects three times as large^2^. We show here that this performance results from a previously unknown type of wing motion. Our experiment combines three-dimensional reconstructions of morphology and kinematics in one of the smallest insects, *Paratuposa placentis* (body length 395 μm). The flapping bristled wing follows a pronounced figure-eight loop that consists of subperpendicular up and down strokes followed by claps at stroke reversals, above and below the body. Computational analyses suggest a functional decomposition of the flapping cycle in two power half strokes producing a large upward force and two down-dragging recovery half strokes. In contrast to heavier membranous wings, the motion of bristled wings of the same size requires little inertial power. Muscle mechanical power requirements thus remain positive throughout the wing beat cycle, making elastic energy storage obsolete. This novel flight style evolved during miniaturization may compensate for costs associated with air viscosity and helps explain how extremely small insects preserved superb aerial performance during miniaturization. Incorporating this flight style in artificial flappers is a challenge for designers of micro aerial vehicles.

---

Driven by curiosity for the smallest objects, scientific exploration of the microscopic world has facilitated the miniaturization of various industrial products. But miniaturization is more than a human-made artifice: inspiring success stories of it are abundant in the living world. For over 300 million years, ecological pressures have forced many insects to become ever smaller. The smallest flying animals have diminished in the course of evolution to become only about 200 μm long^3^.

The physical properties of an object differ in their dependence on its size. Consequently, effects small at the macro-scale can become dominant at the micro-scale, and vice versa^4^. This competition of scaling laws applies also to objects that move through air: in smaller objects, inertia of the air parcels loses its dominance over viscous friction.

Larger animals are generally faster than smaller ones^1^. Nevertheless, some tiniest insects fly surprisingly well. It was revealed recently that minute featherwing beetles (Coleoptera: Staphylinoidea: Ptiliidae) typically fly at the same speeds and accelerations as their relatives Staphylinidae, despite the threefold difference in the body length. Moreover, ptiliids can accelerate twice as fast as larger related staphylinoids^2^. Since the volume of the flight muscles relative to the body volume is the same in Ptiliidae and in larger beetles^5^, the excellent flight performance of the former may be due to the peculiar structure of their wings and the style of using them in flight. Ptiliids have feather-like bristled wings — a condition known as ptiloptery (Fig. 1c). This is arguably the most visually striking modification in the flight apparatus convergently evolved by the smallest representatives of several orders of insects. But the functional benefits of ptiloptery remain largely unknown.

**Fig. 1.**
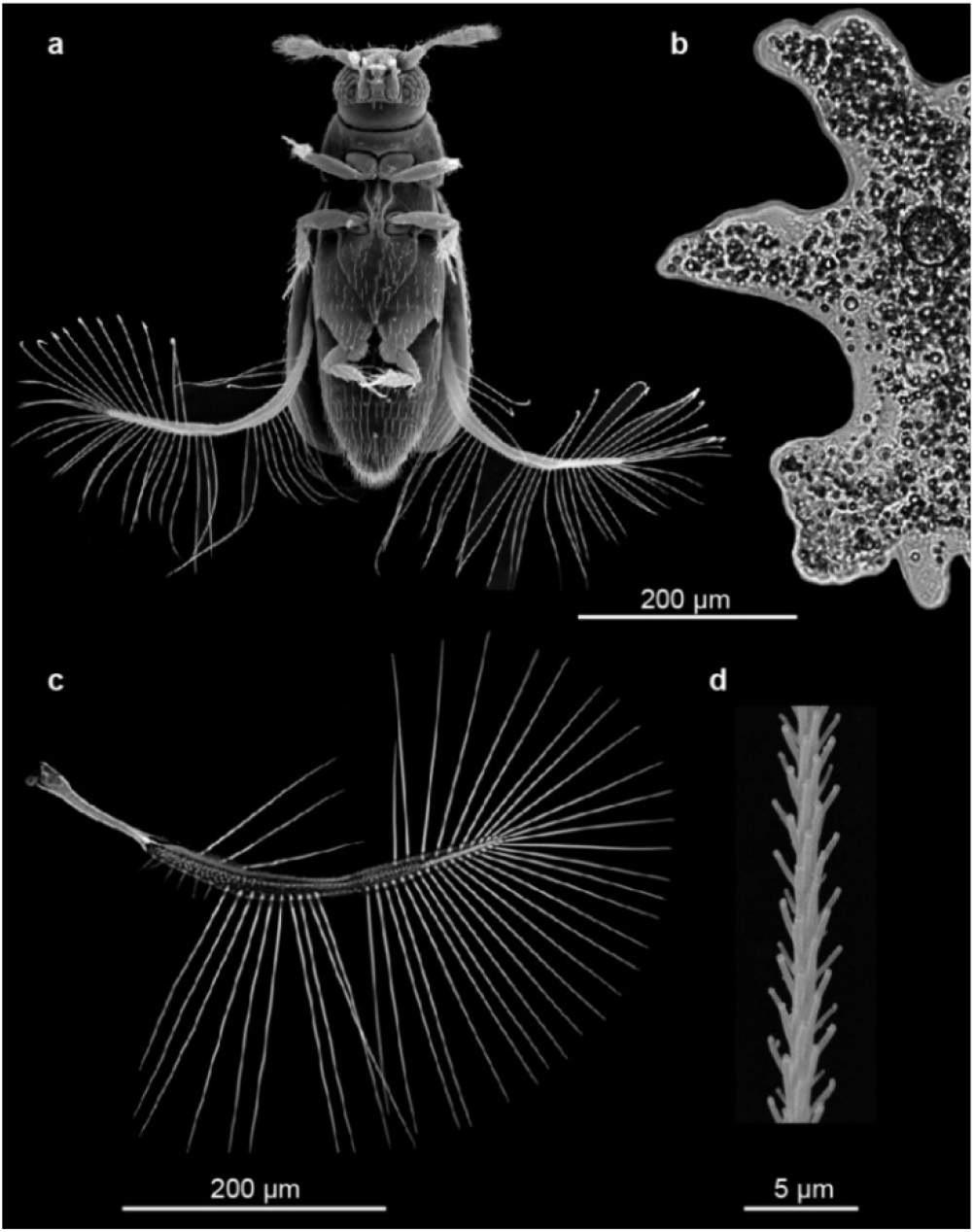
External morphology of *Paratuposa placentis*, SEM images. **a, b,** Body size comparison of *P. placentis* and *Amoeba proteus*. **c,** Wing of *P. placentis*. **d,** Part of seta.

The problem of the flight of minute insects was repeatedly addressed^6,7^, but the experimental data that elucidate it are very limited and have been obtained only for insects that are not at all among the smallest^8,9,10,11^. Although the unsteady aerodynamics of millimetre-size insects such as fruit flies^12,13^ and mosquitoes^14^ received considerable attention in the past decades, studies focusing on the tiny insects remain scarce. Two-dimensional numerical simulations of flows past evenly spaced cylinder lattices showed that the throughflow in the gaps reduces the aerodynamic forces^15,16^. Experiments with mechanical comb-like models suggested a somewhat greater lift to drag ratio during the clap-and-fling phase of the wing motion^17,18,19^, but did not cover the full flapping cycle. Meanwhile, with the state-of-the-art high-speed videography, it became clear that the smaller insects use a wing beat cycle different from that of the larger ones^10,11^, but the role of ptiloptery in this cycle was not considered.

In this study, we analyse the flight of the miniature featherwing beetle *Paratuposa placentis*. We constructed a morphological model using a confocal laser scanning microscope, a kinematical model using synchronized high-speed videography, and a dynamical model using computational methods of solid and fluid mechanics. The combination of these methods offers a comprehensive view on how bristled wings work, and explains why common sub-millimetre flying insects have bristled rather than membranous wings.

## Morphology

*P. placentis* is one of the smallest nonparasitic insects, its body length is 395 ± 21 μm (Fig 1a; hereinafter M ± SD), which is comparable to those of some unicellular protists, such as *Amoeba proteus* (Fig 1b). The body mass of *P. placentis* is 2.43 ± 0.19 μg. The bristled wing consists of a petiole, а narrow wing blade and а fringe of setae (bristles) covered with secondary outgrowths (Fig 1c,d). The wing length is 493 ± 18 μm; the setae occupy 95.1 ± 0.3% of the wing area.

## Kinematics description

We consider relatively slow normal flight with horizontal velocity 0.057 ± 0.014 m/s and 0.039 ± 0.031 m/s vertical velocity (PP2, PP4, PP5, PP12). Wing deformations in flight are insignificant (Supplementary Information), so we neglected them when reconstructing the kinematics. The stroke cycle of *P. placentis* consists of two power strokes, i.e., stages of the cycle during which most force is generated^20^, and two recovery strokes, which represent a considerably modified clap-and-fling pattern (Fig. 2a,c; Supplementary Video 1-5). The morphological downstroke and upstroke are remarkably similar: the angle of attack (AoA) reaches 72.7° during downstroke and 85.0° during upstroke (Fig. 2f), and the pitch angle varies over a wide range, from about 20° to about 180° (Fig. 2b,d). The average wingbeat frequency *f* is 177 ± 16 Hz. Cycle averaged Re, based on mean speed of the radius of gyration, is 9.3 and reaches 19.6 during power strokes, when wing speed is the highest. During the bottom recovery stroke, wings do not clap tightly, the mean (in a series of wing cycles) minimum distance between wing blade tips is 162 ± 94 μm. The setal fringes of the left and right wings can intersect each other early in the fling phases of the recovery strokes. The following descriptions of kinematics and aerodynamics, as well as the illustrations, refer to results obtained for individual PP2. For the results obtained for other specimens, see Supplementary Information.

**Fig. 2.**
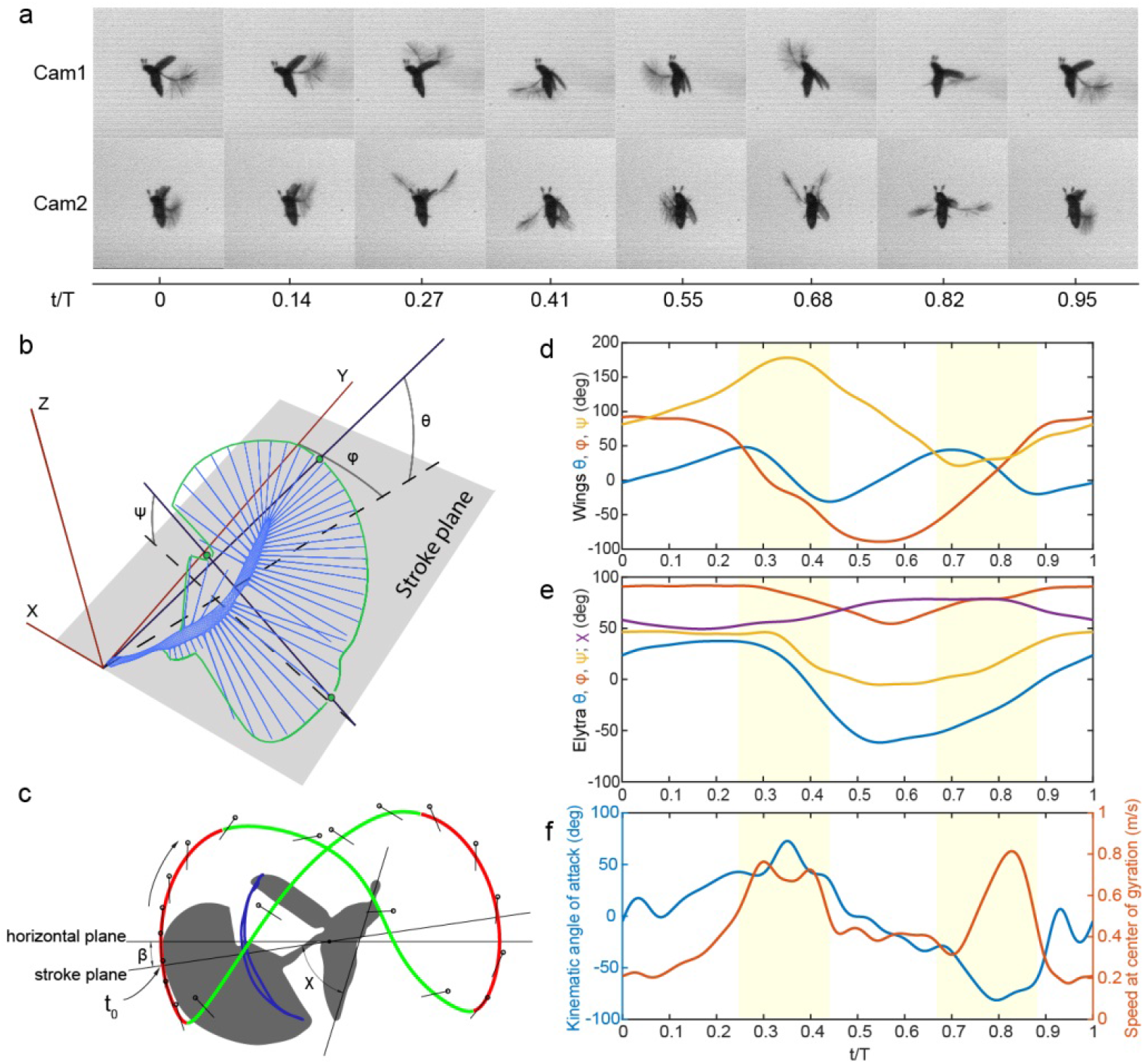
Kinematics of *P. placentis*. **a,** Frame sequence of single stroke in two projections. **b**, Measurement scheme for Euler angles. **c**, Trajectory of wing tip: recovery strokes (magenta line) and power strokes (green line) and measurement scheme for angle of body pitch (*χ*) and pitch of stroke plane to the horizon (*β*). **d**, Wing Euler angles as functions of dimensionless time *t*/*T*, *T* = 1/*f*: stroke deviation (*θ*), positional (*φ*), pitch (*ψ*). **e**, Elytron Euler angles (*θ*, *φ, ψ*); body pitch angle (*χ*). **f**, AoA and wing speed at centre of gyration vs dimensionless time t/T.

## Using aerodynamic lift and drag of the wings for net vertical force generation

High velocity and AoA during the power strokes provide for the necessary weight-supporting time-averaged vertical force of 2.71 μgf (Fig. 3g). For comparison, the estimated mass of the beetle is 2.43 μg and its vertical acceleration is 1.02 m/s^2^. The net contribution of the body and elytra to the vertical force is negligible (Fig. 3g); therefore, we focus on the wings.

**Fig. 3.**
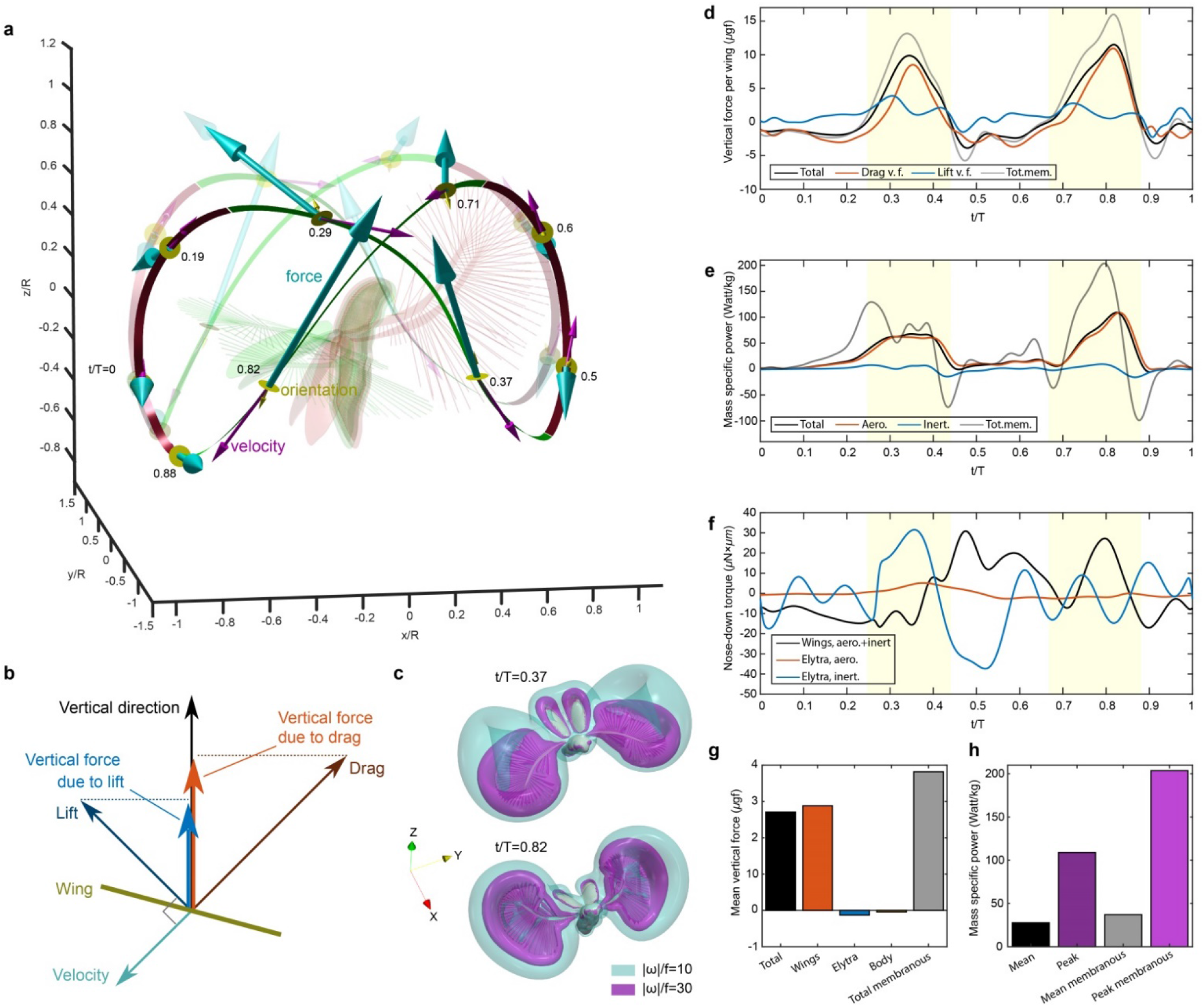
Aerodynamic forces acting on the wings of *P. placentis*. **a,** Wing tip trajectories coloured according to direction of total vertical force: green is down (recovery stroke), red is up (power stroke). Posture at *t*/*T* = 0.6 is shown in red, at *t*/*T* = 0.82 in green. Cyan arrows show aerodynamic force; magenta arrows show wing-tip velocity, yellow disks and arrows show dorsal surface orientation of the wing at nine labelled time instants. Opaque and transparent lines and arrows correspond to right and left wing, respectively. **b,** Vector scheme of forces acting on wing. **c,** Air flow visualization using iso-surfaces of vorticity magnitude (also see Supplementary Video 5). **d,** Vertical aerodynamic force exerted on one wing, vs time. Yellow highlighted zones denote the time span of power strokes. **e,** Body mass-specific aerodynamic and inertial power, and their sum. **f,** Pitching torque about centre of mass. Positive direction is nose down. **g,** Contribution of different parts to total aerodynamic force acting on the beetle in the vertical direction, averaged over the wing beat cycle. **h,** Mean and peak body mass-specific aerodynamic power in computations for bristled and membranous wings.

The wing tips follow wide rounded self-intersecting paths, Fig. 2c, 3a. One cycle consists of two power strokes, when the wings move down, and two recovery strokes, when they move up. They move faster during power stroke than during recovery. Тhe time course of the wing speed (Fig. 2f) shows peaks at *t*/*T* = 0.3 and 0.83. The force and the velocity are anti-aligned (Fig. 3a); their peaks are synchronized (Fig. 3d, 2f). The air flow visualization in Fig. 3c and Supplementary Video 5 reveals a pair of strong vortex rings symptomatic of drag-producing bodies.

The dynamically changing orientation of the wing helps amplify the asymmetry between the power and recovery strokes. Upon each power stroke, the kinematic AoA becomes large simultaneously with the velocity (see Fig. 2f). Thus, the wing produces a great upward force as it quickly moves flat-on with net downward displacement, and produces a small downward force while slowly moving edge-on upwards. The near clap reduces further the parasite downward force upon recovery^21^.

Decomposition of the vertical force exerted on the wing in drag and lift (defined in Methods) is shown in Fig. 3d. The vertical force due to drag has greater positive peaks than that due to lift, but also wider negative valleys. Time averaging yields the mean vertical force breakdown into drag and lift as 32% and 68% of the total, respectively, indicating that the beetle benefits from both components.

## Stabilizing effect of the elytra

The forces are small during the recovery strokes, but the moment arm relative to the centre of mass is great, resulting in a pitching moment great enough to overturn the body around its lateral axis. To compensate, in sync with wing flapping, the insect opens and closes the elytra. Their amplitude (*ψ_max_* - *ψ_min_* = 52°) is greater than in other previously studied beetles: 20° in *Allomyrina dichotoma*^23^, 31.3° in *Trypoxylus dichotomus*^24^ and 45° in *Priacma serrata*^25^. Figs. 2e and 3f explain how the elytra act as inertial brakes. Between t/T = 0 and 0.3, the wings are raised dorsally and produce a nose-up torque. As soon as they spread laterally, the elytra start to close, causing a nose-down recoil torque on the body. As the wings clap ventrally and switch to nose-down torque production, the elytra decelerate and reopen. We estimate that the body pitching oscillation amplitude is halved due to the elytra flapping (for details, see Supplementary Information).

## Comparison with larger beetles and tiny insects of other orders

The large deviation angle amplitude of 85° (Fig. 2b,d) is in striking contrast with the almost negligible wing stroke deviation in some larger beetles (e.g., 9° in *Trypoxylus dichotomus*^24^ and 16° in *Batocera rufomaculata*^26^). Small stroke deviation is typical of the lift-based normal hovering, when the wings move essentially within a plane perpendicular to the direction of the net aerodynamic force. Large deviation is symptomatic of drag-based flight, and it has been observed also in the smallest species among different families of wasps, midges and thrips^10,11^ (Fig. 4a), although *P. placentis* shows an extreme and unique kinematic cycle with two fast power strokes and two slow recovery strokes. The time course of the vertical aerodynamic force showing two narrow positive peaks and two shallow negative valleys has more in common with smaller representatives of other families^10,11^ that with larger beetles^24^. A corollary to the localization of the vertical force in relatively short pulses is that the reaction loads exerted on the skeletomuscular system show unusually high peaks above the mean value. Our calculations suggest that ptiloptery is an efficient solution to the problem of reducing the inertial component of those peak loads. Thus, *P. placentis* can afford to have large enough wings for excellent flight^2^. As Fig. 4b shows, Pliliidae tend to have larger wings relative to their body length than other Staphylinoidea (two-samples Wilcoxon test p = 0.000143).

**Fig. 4.**
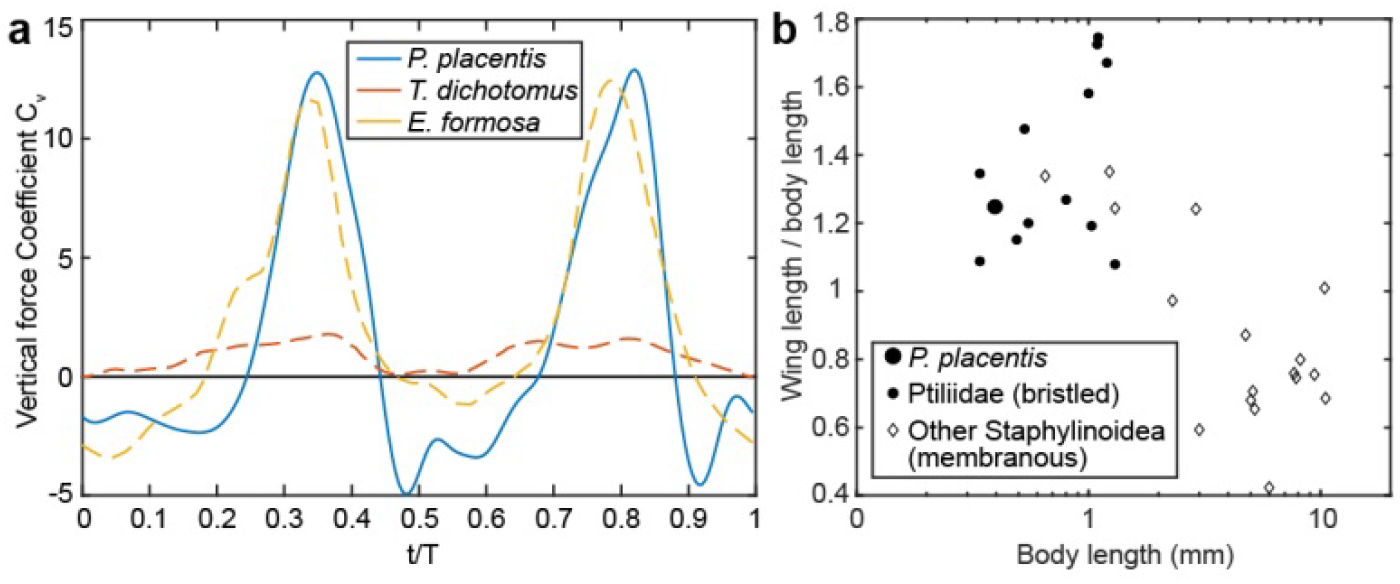
Comparison between *Paratuposa placentis* and other insects. **a,** Time variation of the vertical force coefficient in three different species. Data for the large rhinoceros beetle *Trypoxylus dichotomus*^24^ and for the tiny chalcid wasp *Encarsia formosa*^11^ are adapted, so that the cycle begins with downstroke. The force coefficient is defined as C_V_ = 2F_V_/ρ(2Φr_g_f)^2^S, where F_V_ is vertical force, ρ is density of air, Φ is flapping amplitude, r_g_ is wing geometric radius of gyration, f is flapping frequency, and S is wing area. Note that the bristled wings of *Encarsia formosa*^11^ were modelled as impermeable solid plates. **b,** Wing length relative to body length^27,28,29,30^.

## Ptiloptery and the original flying style help beetles to fly faster

Let us put together four important observations. (I) We have found that bristled wings are lighter than the equivalent membranous wings and have smaller moments of inertia. (II) The inertial forces are proportional to acceleration, i.e., to flapping frequency squared. The inertial power therefore varies as frequency cubed. (III) Since the Reynolds number is relatively low, the aerodynamic force of a wing is proportional to its velocity raised to some exponent γ < 2. The velocity is proportional to the flapping frequency. The instantaneous aerodynamic power, defined as a product of the force and velocity, is proportional to the frequency raised to the exponent 1 + γ < 3. (IV) The wings clap dorsally and ventrally even at low speeds (Supplementary Information). At high speeds, unable to further increase the flapping amplitude, the beetle unavoidably increases the flapping frequency to produce enough aerodynamic thrust to counter the body drag, keeping the flapping amplitude constant. Consequently, the peaks of the inertial power increase, and they do so faster than the aerodynamic power. However, the flapping frequency cannot be increased beyond the point when the muscles become unable to sufficiently accelerate the wing upon power stroke, even if the mean aerodynamic power is low. We arrive at the following conclusion: for a beetle as miniature as *P. placentis* with the cycle-averaged Re as low as 9.3, low wing moment of inertia is a prerequisite to fast flight. This is achieved by ptiloptery.

## Methods

### Material

Adults of featherwing beetles *Paratuposa placentis* (Coleoptera: Ptiliidae) were collected in Cát Tiên National Park, Vietnam, in November 2017. The beetles were collected and delivered to the laboratory together with the substrate for their safety. High-speed video recordings were made on the same day during a few hours after collecting.

### Morphology and morphometry

The material for morphological studies was fixed in alcoholic Bouin solution or in 70% ethanol. Wing structure was studied using a scanning electron microscope (SEM Jeol JSM-6380 and FEI Inspect F50), after dehydration of the samples and critical point drying, followed by gold sputtering. A confocal microscope (Olympus FV10i-O) and a transmitted light microscope (Olympus BX43) were also used, for which the samples were clarified and microscopic slides were made^27^ (Supplementary Information). Measurements were taken from digital photographs in Autodesk AutoCAD software at ten replications (unless otherwise noted). Body weights and weights of particular body parts were calculated based on three-dimensional reconstructions (Supplementary Information).

### High-speed recording

Flight of the beetles was recorded in closed 20 × 20 × 20 mm chambers, custom made of 1.0 mm thick microscopic slides and 0.15 mm cover-glass at a natural level of illumination in visible light. There were 20–30 insects in the flight chamber during the recording. For temperature stabilization the flight chamber was chilled by air fan from the outside. The ambient temperature measured by digital thermocouple was 22 – 24 °C; temperature of the flight chamber was 22 – 26 °C.

High-speed video recordings were made using two synchronized Evercam 4000 cameras (Evercam, Russia) with a frequency of 3845 FPS and a shutter speed 20 μs in infrared light (850 nm LED). The high-speed cameras were mounted on optical rails precisely orthogonal to each other and both 0° from the horizon. Two IR LED lights were placed opposite to the cameras and one light above the flight chamber.

### Measurement of kinematics

For analysis, 13 recordings were selected. For four of them (PP2, PP4, PP5, PP12) we reconstructed the kinematics of body parts in four kinematic cycles for each and performed CFD calculations because the flight of these specimens was especially similar to conventional hovering. In CFD analysis with the membranous wing model, we selected PP2, which does not cross the wings while clapping. This case is convenient for comparing the performance of bristled wings with substitute membranous wings, because is it guarantees that the latter do not intersect. The perimeter of the membrane is formed by lines connecting the tips of the bristles (see the previous study^22^ for more information).

Average wingbeat frequency was calculated as the mean of the wingbeat frequency in all recordings. In each recording, the number of frames was counted in several complete kinematic cycles, 104 cycles in total.

For the mathematical description of the kinematics of the wings and elytra we used the Euler angles system^31,32^ (Fig. 2b) based on frame-by-frame reconstruction of the location of the insect’s body parts (wings, elytra and body itself) was performed in Autodesk 3Ds Max. 3D models of body and elytra were received by confocal microscope image stacking and flat wing model was based on light microscopy photos of dissected wings. We used the rigid flat wing model for reconstruction of the kinematics because deformations of the wings are minor (Supplementary Information). First we prepared frame sequences with four full kinematical cycles in each. The frames were then centred and cropped by point between the bases of the wings and then placed as orthogonal projections. Virtual models of body parts were placed into coordinate system with two image planes. Then we manually changed the position and rotated body parts until their orthogonal projections were superimposed on the image planes. For calculating the Euler angles, a coordinate system was created (Fig. 2a). The X0Y plane is a plane parallel to the stroke plane, and intersecting with the base of wing or elytron, which is positioned in the zero point. To determine the position of the stroke plane, we calculated the Major Axis trend line of the wingtips coordinates instead of the linear trend line^32^, because the wingtip trajectory of *P. placentis* forms a wide scatterplot. Stroke deviation angle (θ) and positional angle (φ) were calculated from the coordinates of the base and apex. Pitch angle (ψ) is the angle between the stroke plane and the chord perpendicular to the line between the base and apex. The body pitch angle (χ) is the angle between the stroke plane and longitudinal axis of the body, calculated as the line between the tip of the abdomen and the midpoint between of the last segments of the antennas. Pitch angle (β) of the stroke plane relative to the horizon was also measured.

For flight speed analysis we performed tracking of the centre of the body (middle point between extreme edges of the head and abdomen) in Tracker (Open Source Physics) in both projections and calculated the instantaneous velocity as well as its vertical and horizontal components in each frame. The obtained speed values were filtered by loess fitting in R (stats package). The minimum distance between the wingblade tips during bottom claps was also calculated.

### Computational fluid dynamics

Time intervals of low-speed flight with duration longer than four wing beats were selected. The angles ϕ, θ, ψ of the left wing, right wing, and elytra and the body angle χ were interpolated on a uniform grid with time step size Δt = 2.6 × 10^−6^ s. By solving numerically ϕ(t) = 0 with respect to t, we identified four subsequent wing beat cycles and calculated the average cycle period T and the wing beat frequency f=1/T. We then spline-interpolated the data for each of the four cycles on a grid subdividing the time interval [0,T] with step Δt, calculated phase averages, then calculated the average between the left and right wing. This yielded the plots shown in Fig. 2c,d. Constant forward and upward/downward flight velocity was prescribed using the time average values of the loess-filtered time series.

The computational fluid dynamics analysis is performed using an open-source Navier-Stokes solver WABBIT^33^, which is based on the artificial compressibility method to enforce velocity-pressure coupling, volume penalization method to model the no-slip condition at the solid surfaces, and dynamic grid adaptation using the wavelet coefficients as refinement indicators. The flying insect was represented as an assembly of five rigid solid moving parts: the two elytra and the two wings move relative to the body, and the body oscillates about its lateral axis. The kinematic protocol is described in Supplementary Information. The computational domain is a 12*R* × 12*R* × 12*R* cube, where *R* is the wing length, with volume penalization used in combination with periodic external boundary conditions to enforce the desired far-field velocity^33^. The computational domain was decomposed in nested Cartesian blocks, each containing 25 × 25 × 25 grid points. Blocks were created, removed, and redistributed among parallel computation processes so as to ensure maximum refinement level near the solid boundaries and constant wavelet coefficient thresholding otherwise during the simulations. The numerical simulations started from the quiescent air condition, continued for a time period of two wing beat cycles with a coarse spatial grid resolution of Δ*x_min_* = 0.00781*R* to let the flow develop to its ultimate periodic state, then the spatial discretization size is allowed to reduce to Δ*x_min_* = 0.00098*R* if the wing was bristled or to Δ*x_min_* = 0.00049*R* if it was membranous, and the simulation continued for one more wing beat period to obtain high-resolution results. The air temperature was 25°C in all cases; its density was equal to ρ = 1.197 kg/m^3^ and its kinematic viscosity was ν = 1.54×10^−5^ m^2^/s, the artificial speed of sound was prescribed as *c*_0_ = 30.38*fR* ^22^. The volume penalization as well as other case-specific parameter values are provided in Supplementary Information.

### Decomposition of the aerodynamic force of a wing into lift and drag components

The drag component of the total instantaneous aerodynamic force acting on the wing is defined as its projection on the direction of the wing velocity at the radius of gyration. The lift component is defined as a vector subtraction of the total force and the drag component. The total lift and drag force vectors are projected on the vertical (z) direction to obtain the time courses shown in Fig. 3d.

## Acknowledgements

A.A.P. is grateful to his mentor R.D. Zhantiev for inspiring him to study miniature insects. The authors thank A.K. Tsaturyan for his helpful discussions.The work of S.E.F., A.A.P., P.N.P. and N.A.L. was supported from by the Russian Science Foundation (project no. 19-14-00045, study of morphology high-speed recording and reconstruction of kinematics). This study was performed using the equipment of the shared research facilities of HPC computing resources at Lomonosov Moscow State University (A.A.P., D.K.), TSUBAME3.0 supercomputer at the Tokyo Institute of Technology (D.K.) and HPC resources of IDRIS (T.E., allocation No. A0102A01664 attributed by the Grand Équipement National de Calcul Intensif (GENCI)). The work of D.K. was supported by the JSPS KAKENHI Grant No. JP18K13693. SEM studies were performed using the Shared Research Facility “Electron microscopy in life sciences” at Lomonosov Moscow State University (Unique Equipment “Three-dimensional electron microscopy and spectroscopy”).

## Author Contributions

A.A.P. conceptualized and designed this study; S.E.F., N.A.L and A.A.P. designed the experiment and collected the data; H.L., F.-O.L and R.O. conceptualized the computational analysis; T.E. and D.K. performed the CFD simulations; S.E.F. and D.K. analysed the data; S.E.F., P.N.P., D.K. and A.A.P. wrote the manuscript. All authors edited the manuscript and approved the final version.

## Supplementary information

### Section 1 – Body size measurements

It was not possible to measure *P. placentis* body mass directly, because the total mass of all specimens was less than the lower capacity of the laboratory balance. Therefore *P. placentis* body mass was calculated based on the body mass of the closely related beetle *Primorskiella* sp., which has a similar body size and proportions (Supplementary Fig. 1). A total of 179 *Primorskiella* sp. specimens were immobilised by carbon dioxide and weighed on a Satrogosm МВ210-А laboratory balance (Satrogosm LLC, Russia). Their total weight was 0.82 mg, and therefore the mean weight of one specimen was 4.58 μg. Confocal stacks of 10 specimens of each species were obtained by Olympus FV10i-O (Olympus Corporation, Japan), using autofluorescence in channels of 488 and 559 nm lasers. The samples were depigmented using hydrogen peroxide solution (Dimethyl sulfoxide + 100% EtOH + 30% H_2_O_2_ in proportions 1+3+1, respectively) for 1–5 days at a temperature of 37 °С. Then the samples were dehydrated in ethanol solutions of increasing concentrations (80, 95, 100, and 100%) and clarified in BABB (Benzyl Alcohol + Benzyl Benzoate in proportions 1+2) for 24 hours. Preparations with the samples were made using two cover glasses and FEP ring spacers. Body reconstructions were built from confocal stacks in Bitplane Imaris using Surface function and automatic image segmentation (Supplementary Fig. 1). Volumes of body reconstructions were measured in the statistical module of the Imaris program. The body volume of *Primorskiella* sp. was 5.33 ± 0.16 nL (hereinafter M ± SD); its density was therefore 0.86 ± 0.02 μg/nL. The body volume of *P. placentis* was 2.83 ± 0.22 nL. Assuming that the body density of both species was equal, the body mass of *P. placentis* was 2.43 ± 0.19 μg.

The body length of each living specimen was measured in high-speed recordings as the distance between the anterior edge of the head and the posterior edge of the abdomen. The three-dimensional body length was calculated based on lengths it two orthogonal projections. The values in frame sequences (from 21 to 106) were averaged for each recording. The average body length of all recorded specimens was 395 ± 21 μm. The body length measured by confocal stacks was 375 ± 1 2 μm. Thus, the body length of living specimens was higher than that of fixed ones (Mann–Whitney U test, p < 0.01), probably because of contraction of intersegment abdominal muscles after fixation. In this connection, we calculated the body mass of the recorded beetles taking into account the changes in their body shape.

The body mass of the recorded specimens in kg was estimated using the formula *m_b_* = *ρK_iso_l*_*b*_^3^, where *K_iso_* = 45.95, *ρ* = 0.86 kg/m^3^ and *l*_b_ is the in-flight body length in m.

**Supplementary Figure 1.**
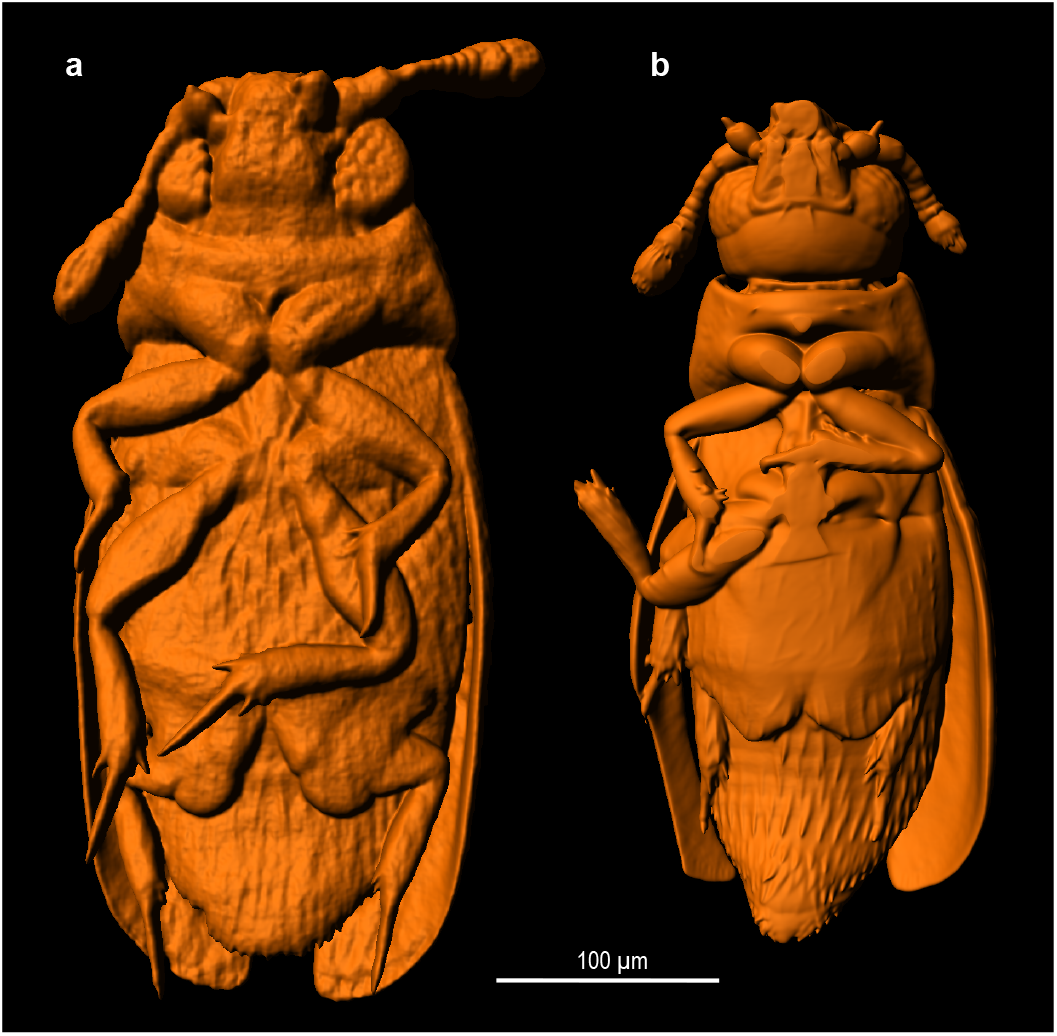
3D reconstruction of featherwing beetles. **a**, *Primorskiella* sp. **b,** *Paratuposa placentis.*

### Section 2 – Wing morphometrics

Ten dissected wing preparations were manufactured according to the protocol described in the previous study^1^. The morphometric characteristics (wing length, total wing area and area of the membranous part plus petiole) were measured in Autodesk AutoCAD, using images taken with a light microscope.

### Section 3 – Wing deformations

For evaluation of transverse wing deformations, we selected frames that showed the wing in lateral view during power strokes: 12 upstroke frames and 10 downstroke frames. Since at these moments the speed and angle of attack and therefore drag are near the local maximum, the wing is under the highest lateral loads and most deformed.

In lateral view three points can be identified on the wing: the base (P_1_), the wingtip (P_5_) and the point on the border between the petiole and the membrane (P_3_). P_3_ was marked at the border that divides the wing into the distal part (membrane plus bristles) and the proximal part (petiole). We marked point P_2_ in the geometric middle of petiole, and point P_4_ in the geometric middle of the distal part (Supplementary Fig. 2b, c).

**Supplementary Figure 2.**
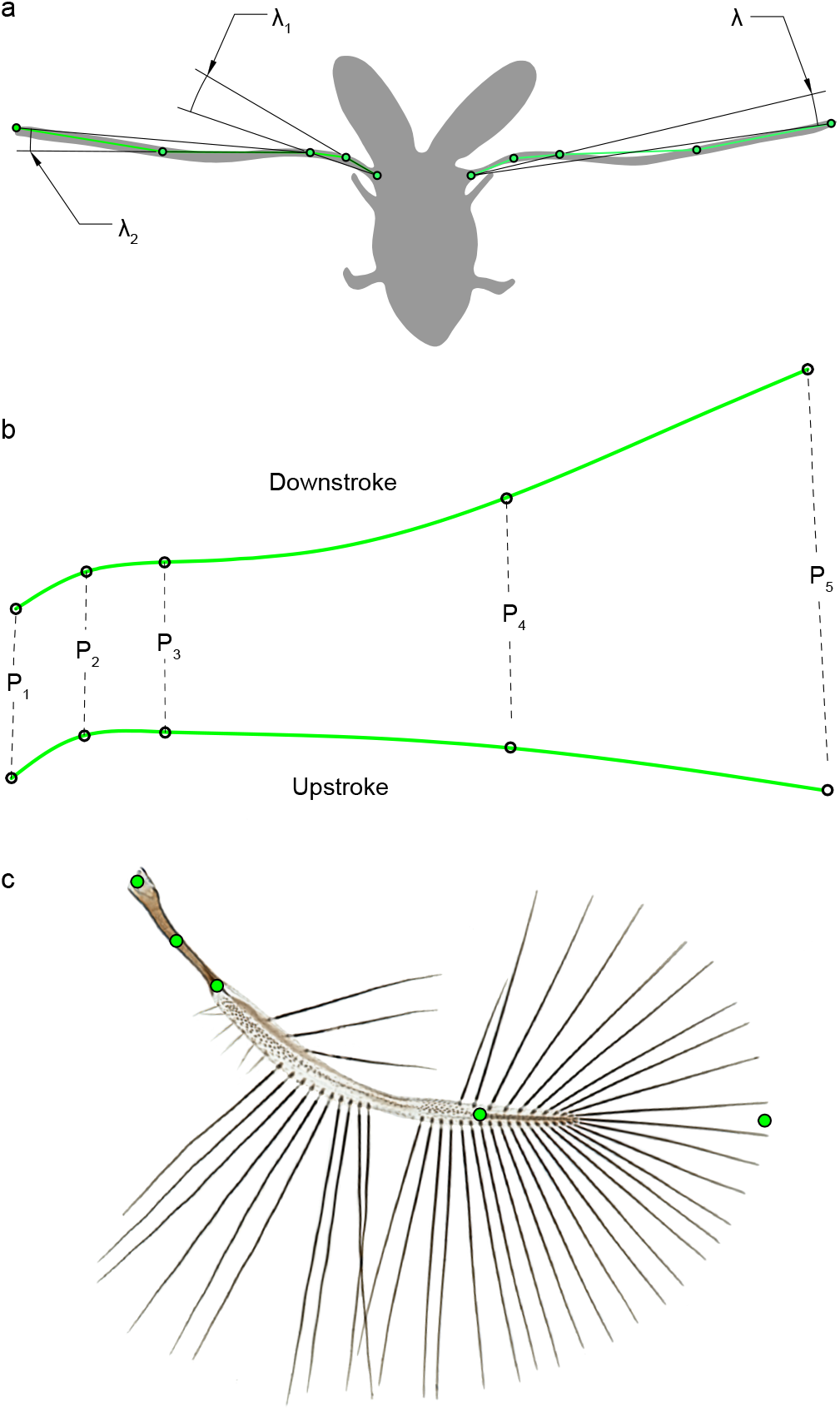
Deformations of the longitudinal wing section during downstroke and upstroke. **a,** Measurement of deformation angles of wing (λ), petiole (λ_1_) and distal part of wing (λ_2_). **b,** Averaged shape of longitudinal wing section during downstroke and upstroke; location of landmarks P_1_–P_5_, lateral view. **c,**Location of marks P_1_–P_5_, dorsal view.

The curvature of the petiole was measured as the angle λ_1_ between the sections P_1_P_2_ and P_1_P_3_. The curvature of the distal part is the angle between P_3_P_4_ and P_3_P_5_. The total wing curvature is the angle λ between P_1_P_3_ and P_1_P_5_. The deformation range of the wing is the sum of the average upstroke and downstroke λ.

The deformation range of the distal part of the wing is 7.6°; the petiole deforms to a smaller degree, on average within 3.5° (Supplementary Table 1). The average deformation range of the wing is 16.5°. The wing has an S-shaped profile during the downstroke and a J-shaped profile during the upstroke (Supplementary Fig. 2b). Resolution did not allow us to measure deformations of the bristles separately, because the bases of the bristles could not be identified. But it was visible that the curvature between P_4_P_5_ was less than between P_3_P_4_, so the deformation of the bristles was smaller than deformations of the membranous part of the wing.

**Supplementary Table 1.** Mean values of wing deformation angles during downstroke and upstroke.

| Deformation angle | $\lambda_1$, ($^\circ$) | | $\lambda_2$, ($^\circ$) | | $\lambda$, ($^\circ$) | |
| --- | --- | --- | --- | --- | --- | --- |
|  | Mean | SD | Mean | SD | Mean | SD |
| Upstroke | 14.0 | 2.7 | 2.4 | 1.0 | 18.2 | 5.7 |
| Downstroke | 10.5 | 2.1 | 5.2 | 2.6 | 1.7 | 7.4 |
| Range | 3.5 |  | 7.6 |  | 16.5 |  |

Wing drag makes up most of the resulting aerodynamic force and is proportional to the projection of the wing onto the plane perpendicular to its instantaneous velocity. Because the area and velocity of the petiole are minor, its contribution to drag can be neglected. The area of the projection and the drag of the rest of the wing depend on the curvature of the distal part and are proportional to cosλ_2_, which varies within 1% of the estimated resting position. Such a small influence on the aerodynamics allows us to ignore deformations in kinematics reconstructions and CFD calculations and use rigid wing models.

### Section 4 – 3D reconstruction of kinematics

The time courses of the Euler angles of the wings and elytra and body pitch angle of individual beetles PP2, PP4, PP5 and PP12 is shown in Supplementary Fig. 3a. There are minor differences between individuals. Vertical and horizontal components of instantaneous acceleration are strongly varying depending on stage of the cycle (Supplementary Fig. 3b). Vertical acceleration synchronised with time course of the lift force.

The wings clap dorsally and near-clap ventrally. During ventral near-claps after downstroke, the minimal distance between tips of the membranous parts of the wings of beetles PP2, PP3, PP5, PP6, PP8, PP10 and PP12 is 0.40±0.23 body lengths. It correlates negatively with the vertical component of the instantaneous body velocity (Spearman r_s_ = −0.78, p < 0.05, n = 25). It can be assumed that with increasing vertical velocity, viscous drag force decreases the lift, and the insect compensates this drag through denser wing clapping.

**Supplementary Figure 3.**
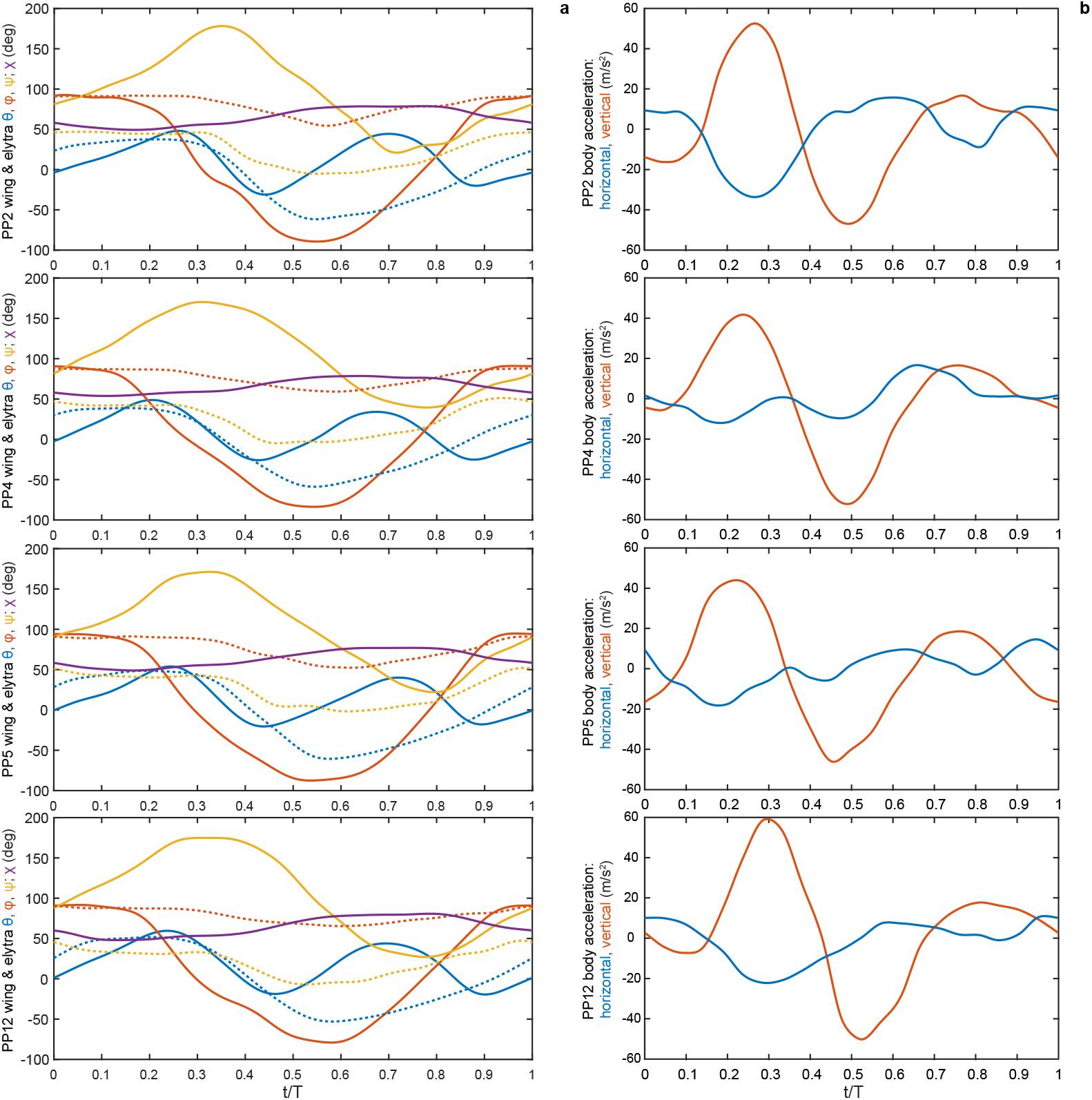
Kinematics description of PP2, PP4, PP5, PP12. **a,** Euler angles of wings and elytra and body pitch angle. **b**, Vertical and horizontal components of body acceleration.

### Section 5 – Building 3D models for CFD

The insect is represented in our 3D models as an assembly of five rigid solid parts: two elytra, two wings, and a body. Supplementary Fig. 4 shows the bristled model of the left wing. It is similar to the model used in the earlier rotating wing simulations^2^. The main difference is in the out-of-plane deviation, which is substantial in the proximal part of the blade, according to the confocal microscope data. The distal part of the wing is essentially planar. The wing length is denoted as *R*. In the aerodynamic calculation, the bristles are modelled as circular cylinders with diameter *b* = 0.00388*R*, which is justified by the prior work^2^. The central blade thickness is *h_b_* = 0.008*R* according to the confocal microscope data. The right wing used in the simulations differs from the left wing by a small rotation of the bristles in the *x_w_*-*y_w_* plane around their respective base points, so that the bristle tips deflect by a distance four times as large as the bristle diameter *b*. This is sufficient to prevent collision between the wings when they clap.

Supplementary Fig. 5 shows the membranous wing model. It is constructed by sealing the gaps between the bristles. This is similar to gluing adhesive tape sheets in prior experiments^2^. To simplify the geometrical processing, the membranous wing is made flat. It is only used in conjunction with the wing kinematics of individual number 2. The latter is such that the wings do not intersect when they clap. In the aerodynamic calculation, the wing thickness at any point is equal to the blade thickness *h_b_*. This enables computing with the minimum discretisation step Δ*x_min_* twice as large as in the bristled wing aerodynamic simulations. The left and the right membranous wings are mirror images of each other.

**Supplementary Figure 4.**
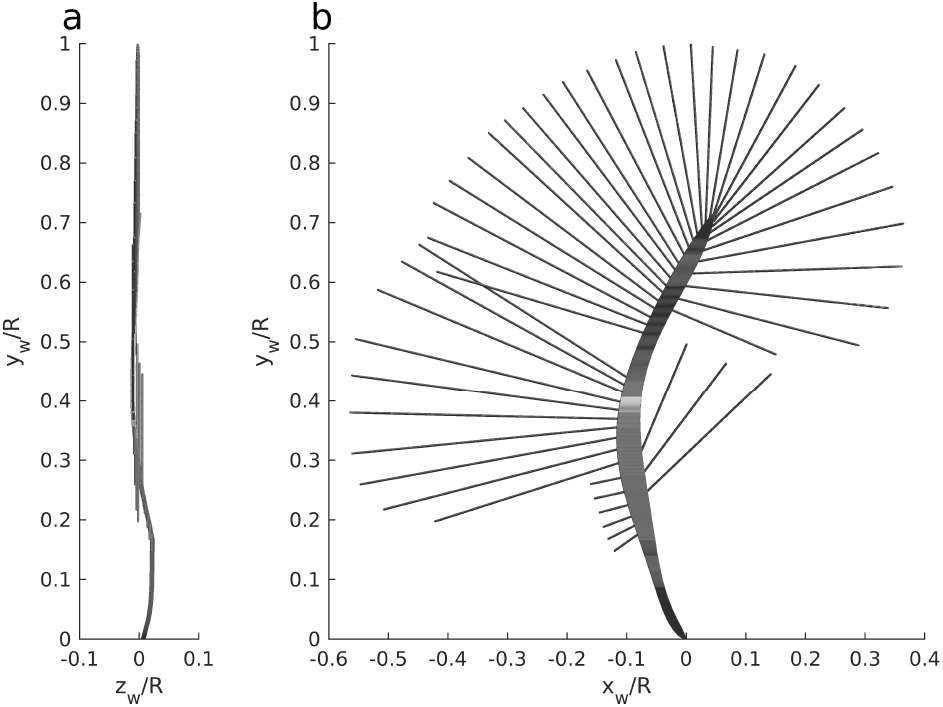
Orthographic projection view of the bristled wing model (left wing). Distances are scaled by the wing length *R*. **a,** View from the trailing edge side. **b,** Morphologically dorsal view.

**Supplementary Figure 5.**
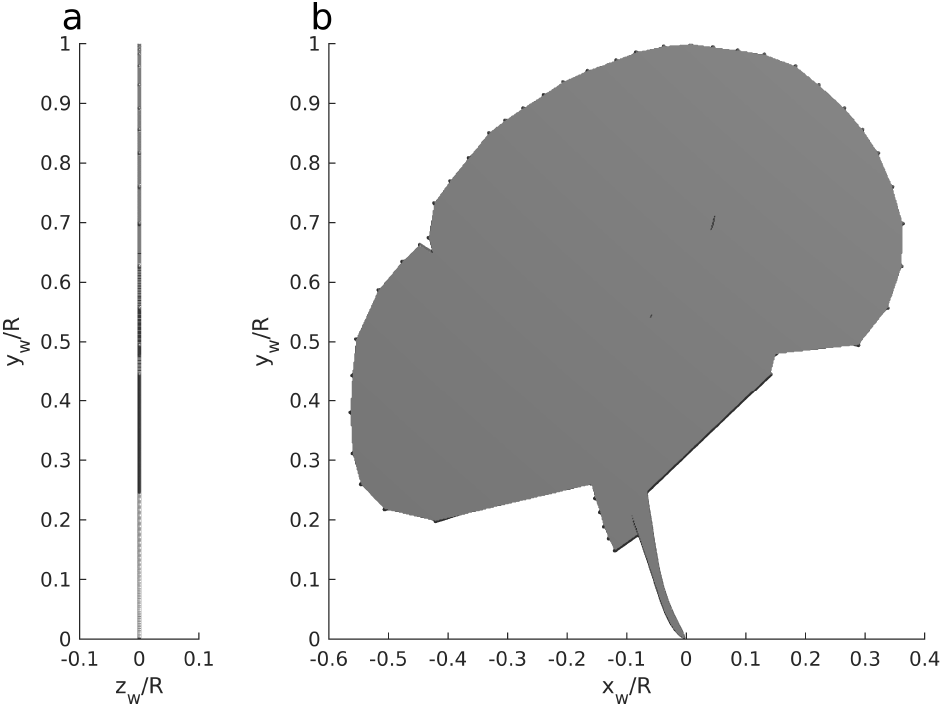
Orthographic projection view of the membranous wing model. Distances are scaled by the wing length *R*. **a,** View from the trailing edge side. **b,** Morphologically dorsal view.

The two elytra are also mirror images of each other. The volumetric models are based on confocal microscope data (Supplementary Fig. 6). However, the proximal part has been modified to avoid intersection with the body. For the elytra and for the wings alike, the centre of rotation is at the origin of the respective local coordinate system. The thickness in the aerodynamic calculation is uniform, *h_e_* = 0.008*R*. Note that the assumed uniform thickness of the elytra and the wings in the aerodynamic geometrical models described here is, in general, different from the thickness in the inertia calculations.

**Supplementary Figure 6.**
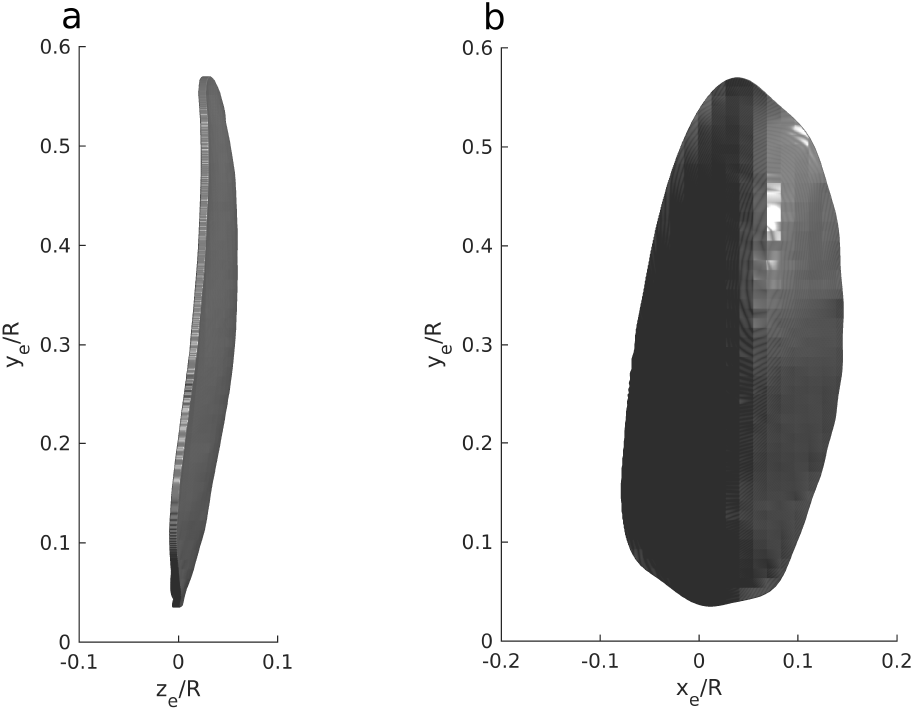
Orthographic projection view of the left elytron model. Distances are scaled by the wing length *R*. **a,** View from the trailing edge **b,** Morphologically dorsal view.

The body model (Supplementary Fig. 7) is also based on information from the confocal microscope, and it is stored as a triangulated surface. Note that it is not perfectly bilaterally symmetric. It is also noteworthy that, while the position of the hinge points in the model closely matches the attachment location of the wings measured from SEM images, the position of the elytron hinges has been slightly offset to prevent intersection between the elytra and the wings as they undergo rigid solid body rotation.

**Supplementary Figure 7.**
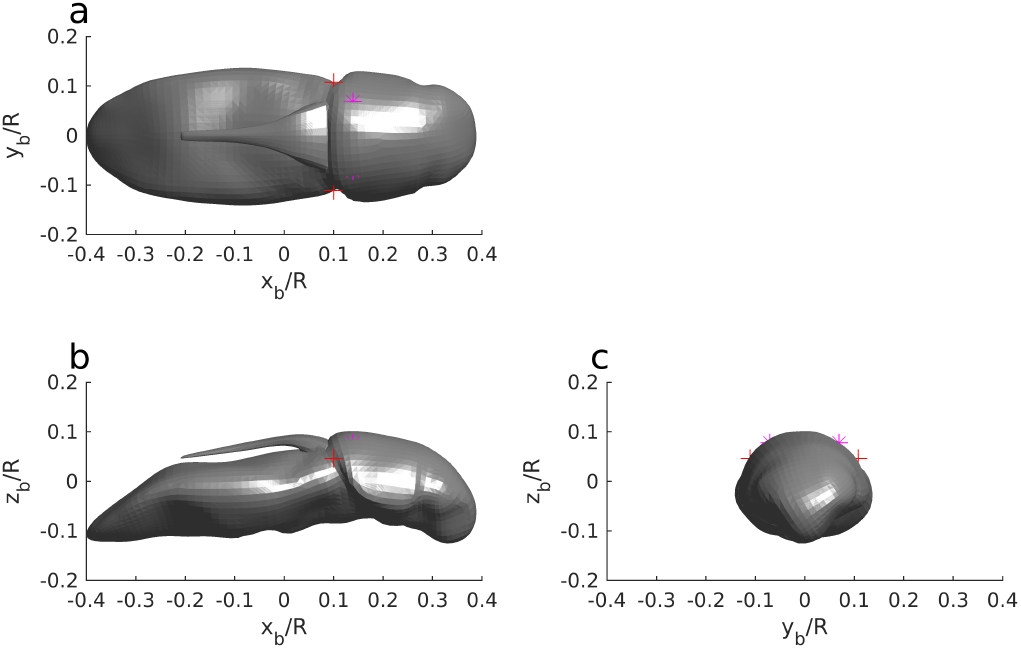
Orthographic projection views of the body model. The red “+” signs show the wing hinge points. The magenta asterisks show the virtual hinge points of the elytron. Distances are scaled by the wing length *R*. **a**, Dorsal view. **b**, Lateral view. **b**, Front view.

### Section 6 – Calculations of moments of inertia for body, wing and elytron

The second moment of inertia of the body about the pitching axis was calculated in Autodesk Inventor Professional 2019 (Autodesk Inc., U.S.A.) using the CAD model of the body shown in Supplementary Fig. 7. The surface model described above, after geometric simplification, was “converted to base feature” of Autodesk Inventor and the inertial properties were evaluated assuming uniform density distribution. In combination with the body mass scaling formula, we obtained the following isometric scaling for the moment of inertia in kg·m^2^ as a function of the body length in m: *J_b_* = 2.14*l_b_*^5^.

After appropriate rescaling, we obtained the principal moments of inertia: *I_xx_* = 0.30·10^−17^ kg·m^2^ about the anterior-posterior axis, *I_yy_* = 2.06·10^−17^ kg·m^2^ the about the lateral axis and *I_zz_* = 2.15·10^−17^ kg·m^2^ about the dorsal-ventral axis. The values provided in this section are scaled to the body length 0.37 mm and mass 2.43 μg, which correspond to individual PP2 and coincide with the statistical averages. Moments of inertia of individuals PP4, PP5 and PP12 are calculated by rescaling the data for PP2 in proportion with the body length to the power of 5.

The wing mass and moments of inertia were calculated by summing up the contributions from the central membranous part (blade) and the peripheral setae (bristles). The mass of the blade (0.0162 μg) was evaluated as a product of the volume enclosed by the SEM surface model obtained by confocal microscopy stacking (1.346·10^−5^ mm^3^) and the cuticle density. The latter was taken as 1200 kg/m^3^ in all calculations related to the wings and the elytra^3^. To calculate the mass of the setae, we first estimated their linear density (0.96 μg/m) using a 3D model consisting of a short cylindrical segment and a few secondary outgrowth elements. The same model was used in our previous study^2^ to select the aerodynamically equivalent circular cylinder section. The mass of each seta was then found by multiplying its length by the linear density. By summing up the masses of all setae, we determined the mass of the peripheral bristled part of the wing (0.0073 μg). Thus, the full mass of a wing including the blade and the setae is 0.0235 μg. This constitutes 0.97% of the body mass.

Subsequently, we calculated the moments of inertia with respect to the hinge point which is the centre of solid body rotation of the wing model used in the aerodynamic simulations. We neglected the out-of-plane deviation of the blade and applied a simple rectangle 2D quadrature rule with the discretisation step of 50 μm. The moments of inertia of the individual setae were calculated using the formula for a thin rod at an angle and the parallel axis theorem. The summation of the above contributions yielded the following values for the non-zero components of the bristled wing inertia tensor: *I_xx_* = 1.45·10^−18^ kg·m^2^, *I_yy_* = 0.12·10^−18^ kg·m^2^, *I_zz_* = 1.57·10^−18^ kg·m^2^, *I_xy_* = −0.16·10^−18^ kg·m^2^. Note that the direction of the x-axis is from the trailing edge to the leading edge, y-axis is from the proximal part to the apex, the z-axis is complimentary, and the origin of the coordinate system is at the wing hinge (Supplementary Fig. 4).

We used the same method to evaluate the mass and the moments of inertia of a wing with hypothetical smooth cylindrical setae having equivalent circular cylinder section (i.e., having a larger diameter and producing the same aerodynamic force as the setae with secondary outgrowths). This diameter, b = 1.9 μm, is 2.1 times as large as the inner diameter of the real seta^2^. The full mass of a wing including the blade and the aerodynamically equivalent cylindrical setae is equal to 0.0421 μg. The corresponding components of the inertia tensor are about twice as large as for the original wing: *I_xx_* = 3.37·10^−18^ kg·m^2^, *I_yy_* = 0.36·10^−18^ kg·m^2^, *I_zz_* = 3.73·10^−18^kg·m2, *I_xy_* = −0.37·10^−18^ kg·m^2^.

The mass and the moment of inertia of the equivalent membranous wing have been determined under the assumption that the membrane thickness is equal to the minimal thickness of the membranous part of the wing obtained by of measuring of cross sections of the wing obtained from histological preparations. Thus, we prescribed the equivalent membrane thickness as a constant value of 0.98 μm. We used the above-mentioned 2D quadrature rule to calculate the area enclosed by the outer contour of the blade and the line connecting the tips of the setae^2^, as well as the moments of area. After multiplying by the membrane thickness and the cuticle density, we found that the equivalent membranous wing mass is 0.1637 μg. This is almost 7 times as great as the mass of the bristled wing. The non-zero components of the equivalent membranous wing inertia tensor are *I_xx_* = 15.70 · 10^−18^ kg·m^2^, *I_yy_* = 2.55 · 10^−18^ kg·m^2^, *I_zz_* = 18.25 · 10^−18^ kg·m^2^, *I_xy_* = −1.90 · 10^−18^ kg·m^2^. Note that these values are almost one order of magnitude greater than in the case of the bristled wing.

To evaluate the mass and the moments of inertia of the elytron, we used the same method as for the central membranous part of the wing. The mass was estimated using a 3D model as 0.1740 μg. Then, the moments of inertia were obtained by 2D integration, neglecting the out-of-plane deviation and assuming uniform thickness distribution, yielding *I_xx_* = 7.16 · 10^−18^ kg·m^2^, *I_yy_* = 0.30 · 10^−18^ kg·m^2^, *I_zz_* = 7.46 · 10^−18^ kg·m^2^, and *I_xy_* = 0.82 · 10^−18^ kg·m^2^.

### Section 7 – Kinematic model for CFD

The wings and the elytra rotate about the hinge points fixed in the body reference frame (Supplementary Fig. 7). For the CFD simulation, the angles ϕ, θ, ψ originally measured in the flight experiment are converted into the Euler angles compatible with the CFD solver^4^, relative to the anatomical stroke plane. We first calculated the Euler angles in the same reference frame of the aerodynamic stroke plane as used in the definition of ϕ, θ, ψ: the positional angle is equal to ϕ, the elevation angle is equal to θ, but the feathering angle is defined as the rotation angle about the longitudinal axis of the wing, i.e., it is different from ψ. Denoting the feathering angle as α, and the respective angle between the wing plane and the stroke plane as ψ^†^(α), we use a genetic optimisation algorithm CMA-ES^5^ to find the values of α that minimise the residue |ψ^†^(α) − ψ| and therefore define the same wing orientation as the originally calculated angle ψ. We calculate the phase average and left-right average values of α as described above.

We then use the morphological model of the wing to define marker points. Each seta takes 100 points uniformly distributed along the axis between its root and its tip. A total of 600 points are placed along the central blade midline. Knowing the Euler angles of the wings in the aerodynamic stroke plane reference frame and the distance between the hinge points, it is possible to determine the spatial position of the markers on the flapping wings. We store those instantaneous positions at 201 time instants between *t* = 0 and *t* = *T*. Subsequently, we evaluate the instantaneous orientation of the body at the same time instants using the mean value and the first Fourier mode of the body angle χ. We define the anatomical stroke plane that moves with the body, and coincides with the aerodynamic stroke plane when the body angle is equal to its mean value.

The CFD simulation requires the Euler angles of the wing measured with respect to the anatomical stroke plane. Their values are found using optimisation. We start from a random guess, evaluate the corresponding position of the markers on the wings in 3D, calculate the root-mean-square distance between these markers and those determined earlier in the aerodynamic stroke plane reference frame, and employ the CMA-ES to minimise that distance. The optimisation is performed for the wings and for the elytra. It yields the values of positional, elevation and feathering angles with respect to the anatomical stroke plane that oscillates with the body. The values are available at 201 time instants between *t* = 0 and *t* = *T*. To evaluate the Euler angles at any arbitrary time instant and ensure periodicity, we spline-interpolate on a uniform grid with 1024 points, apply the fast Fourier transform, and store the first M = 11 modes in the form of the sine and cosine coefficients, as required by the CFD solver^4^.

The orientation of the body was prescribed by defining the time evolution of the pitch angle between the horizontal plane in the laboratory reference frame and the longitudinal axis of the body *x_b_*. Note that in this study the body pitch angle is positive when the body is oriented nose up (this is opposite to the sign convention used previously^4^). Fourier analysis was performed and only the mean, the first sine and the first cosine coefficients were retained.

The anatomical stroke plane does not move with respect to the body. It is tilted by an angle η relative to the body transverse plane *y_b_*-*z_b_*, in the notation of work by Engels et al. so that the average orientation of the anatomical stroke plane coincides with the orientation of the original aerodynamic stroke plane^4^.

Supplementary Fig. 8 summarises the time evolution of the Euler angles of the wing and elytron relative to the body, and the body pitch angle, for four individuals, as used in the CFD simulations. The anatomical stroke plane angle η takes the values −33.5, −36.6, −28.5 and −35.8° for the four analyzed individuals, respectively.

**Supplementary Figure 8.**
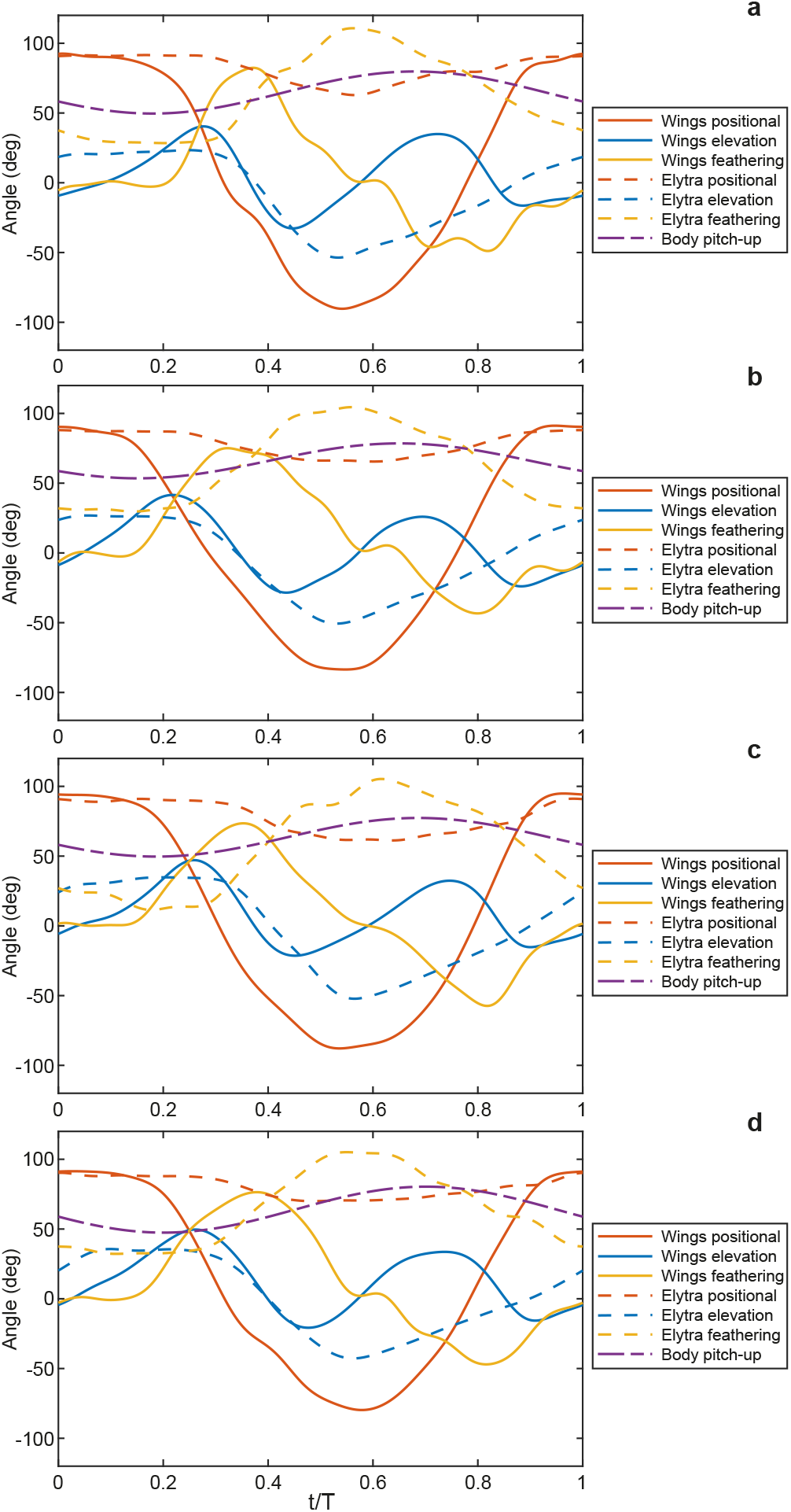
Euler angles of the wings and elytra and the body pitch-up angle between the horizontal plane and the longitudinal axis of the body. **a**, PP2. **b**, PP4. **c**, PP5. **d**, PP12.

### Section 8 – CFD simulation setup

The computational fluid dynamics analysis is performed using an open-source Navier-Stokes solver WABBIT^6^. It is based on the artificial compressibility method to enforce the velocity-pressure coupling, volume penalisation method to model the no-slip condition at the solid surfaces, and dynamic grid adaptation using the wavelet coefficients as refinement indicators. The morphology and kinematics described above define the time-varying position of the volume penalisation mask function and the internal velocity field of the solid parts.

The computational domain is a 12*R* × 12*R* × 12*R* cube (Supplementary Fig. 9), where *R* is the wing length. The volume penalisation is used in combination with periodic external boundary conditions to enforce the desired far-field velocity^6^. The computational domain is decomposed in nested Cartesian blocks, each containing 25 × 25 × 25 grid points. Blocks are created, removed and redistributed among parallel computation processes so as to ensure maximum refinement level near the solid boundaries and constant wavelet coefficient thresholding otherwise during the simulations. Since the flow is far from being turbulent but Δx_*min*_ is small for geometrical reasons, the threshold value for thresholding wavelet coefficients is fixed at ε = 10^−2^. With this choice, the most refined grid blocks are those that contain the solid surface.

**Supplementary Figure 9.**
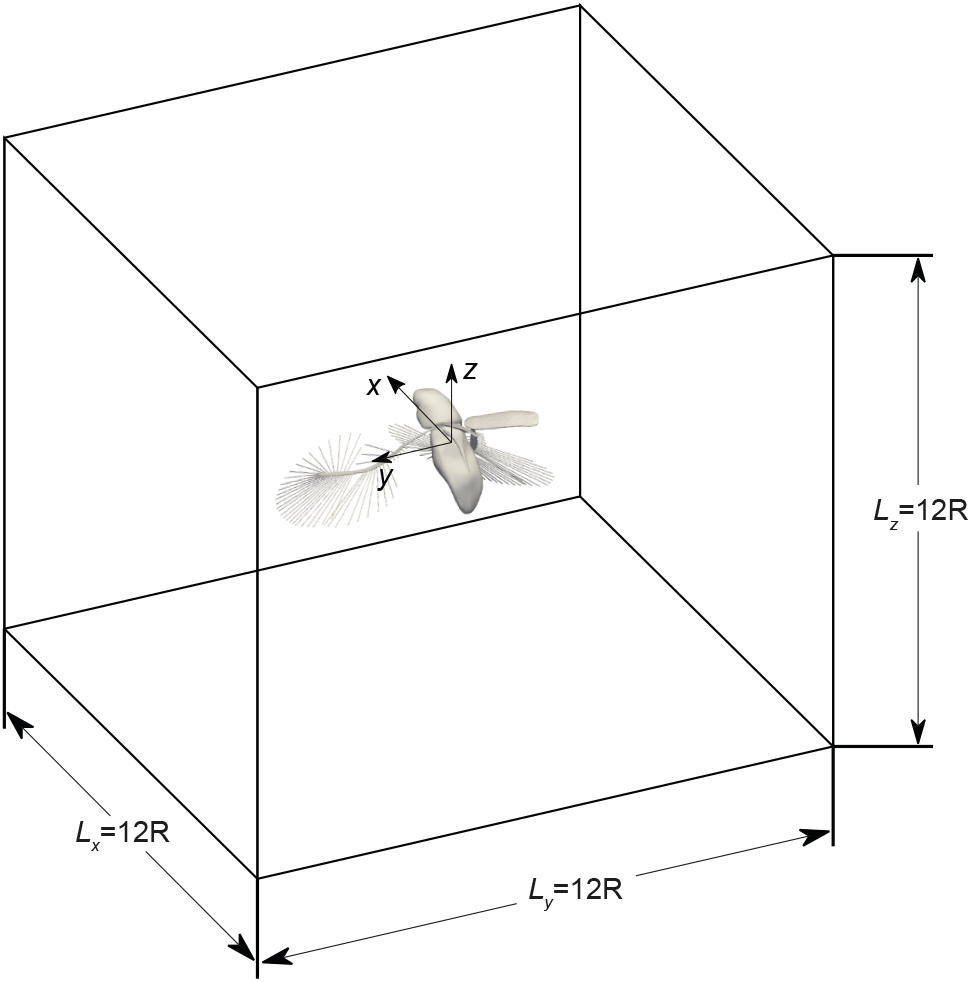
Computational domain.

The numerical simulations start from the quiescent air condition, continue for a time period of two wing beat cycles with a coarse spatial grid resolution of Δ*x_min_* = 0.00781*R* to let the flow develop to its ultimate periodic state, then the spatial discretisation size is allowed to reduce to Δ*x_min_* = 0.00098*R* if the wing is bristled or to Δ*x_min_* = 0.00195*R* if it is membranous, and the simulation continues for one more wing beat period to obtain high-resolution results. The air is at 25 °C temperature in all cases, its density ρ = 1.197 kg/m^3^ and its kinematic viscosity is ν = 1.54·10^−5^ m^2^/s.

All numerical simulations are performed with the artificial speed of sound prescribed as *c*_0_ = 30.38*fR*. This value has been selected based on the preparatory comparison between CFD simulations and dynamically scaled rotating wing model experiments^2^. The values assigned to the volume penalisation parameter vary in proportion with Δ*x_min_*^2^/ν. The high-resolution results are obtained with *C*_η_ = 6.48·10^−7^*f*, 7.50·10^−7^*f*, 7.29·10^−7^*f* or 6.66·10^−7^*f*, depending on the individual number PP2, PP4, PP5 or PP12, respectively.

The low Reynolds number of the flow requires that the step of explicit time integration be proportional to Δ*x_min_*^2^. This constraint is handled by using the Runge–Kutta–Chebyshev time-stepping schemes^7^ with large stability regions designed so as to encompass all eigenvalues of the linear part of the differential operator in each of the four cases. These schemes are second-order accurate, and the number of stages *s* is relatively large to guarantee the desired stability: *s* = 22, 20, 21, 22 for the individuals PP2, PP4, PP5 or PP12, respectively.

### Section 9 – CFD results

The time evolution of the aerodynamic force acting on the beetle is shown in Supplementary Fig. 10. In all cases, the vertical force has two positive peaks during the power strokes, and it is negative during the recovery strokes. The horizontal force also peaks during the two power strokes, but the sign changes from negative to positive. The mean vertical force is positive. The calculated value is large enough to support the weight of the insect, with the small discrepancy which is partly due to the non-zero vertical acceleration of the beetle in the experiment, and partly due to modelling error. The mean horizontal force is small. These mean values are shown in Supplementary Table 2.

Newton’s second law provides a useful relation for diagnosing the accuracy of the aerodynamic force computation. Supplementary Table 2 shows the two components of the residual error in the law of motion evaluated after substituting the computed mean force and the measured acceleration,

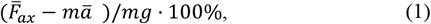

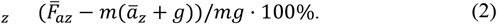

The r.m.s. values of *e_x_* and *e_z_* across the four individuals are, respectively, 9.4% and 12.1%. Note that this error includes the inaccuracy of the CFD modelling and the uncertainty of the body mass evaluation.

**Supplementary Figure 10.**
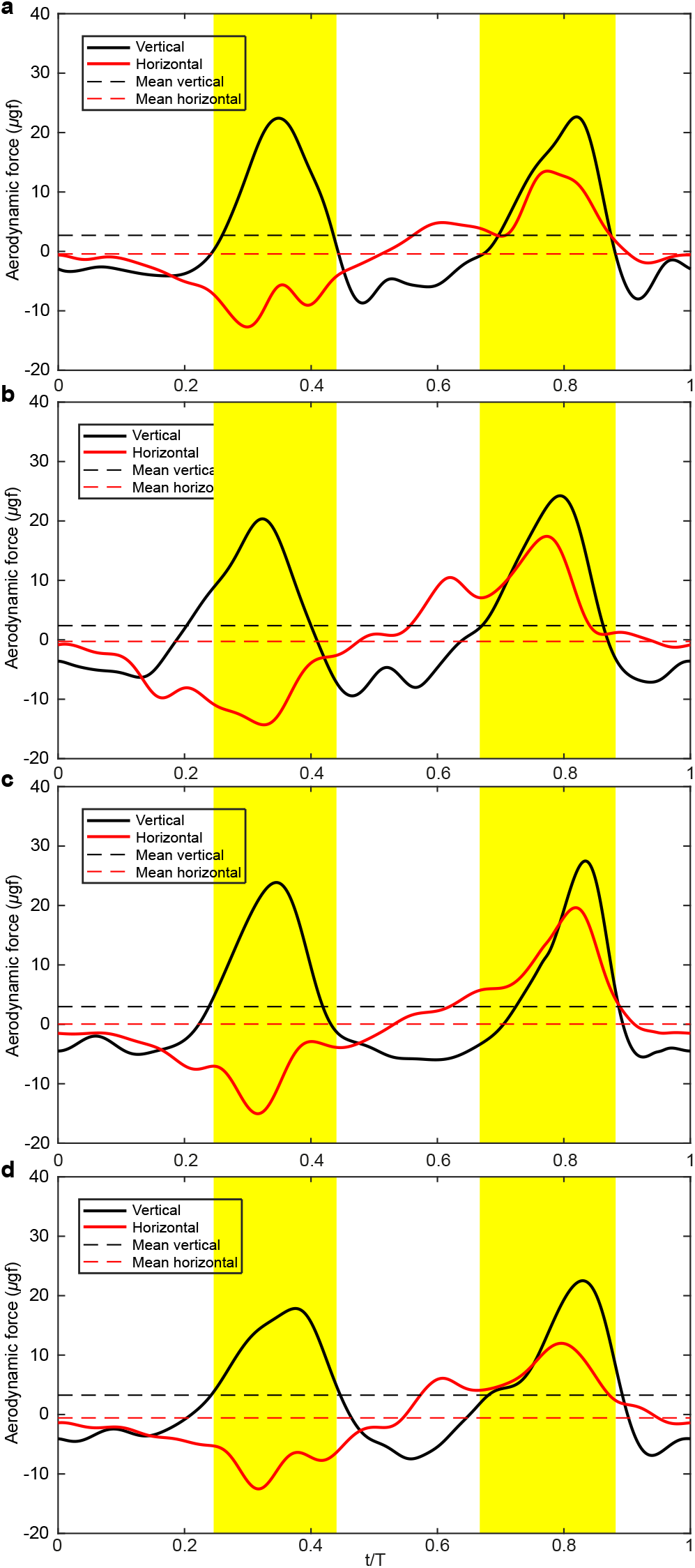
Vertical and horizontal components of the aerodynamic force exerted on the insect. **a**, PP2. **b**, PP4. **c**, PP5. **d**, PP12.

**Supplementary Table 2.** Input and output parameters of the CFD simulations specific to individuals.

| Individual number | PP2 | PP4 | PP5 | PP12 |
| --- | --- | --- | --- | --- |
| Flapping frequency $f$ (Hz) | 171.0 | 190.9 | 190.9 | 183.6 |
| Anatomical positional angle amplitude $\Phi$ ($^{\circ}$) | 182.9 | 174.8 | 182.7 | 170.9 |
| Aerodynamical stroke plane angle $\beta$ ($^{\circ}$) | 8.2 | 12.6 | 2 | 9.7 |
| Anatomical stroke plane angle $\chi$ ($^{\circ}$) | -33.5 | -36.6 | -28.5 | -35.8 |
| In-flight body length $l_b$ (mm) | 0.395 | 0.405 | 0.397 | 0.389 |
| Wing length $R$ (mm) | 0.493 | 0.505 | 0.495 | 0.485 |
| Mean velocity at radius of gyration $R_g$ (m/s) | 0.444 | 0.475 | 0.480 | 0.438 |
| Estimated body mass $m_b$ (μg) | 2.43 | 2.62 | 2.46 | 2.32 |
| Mean horizontal velocity $\bar{V}_x$ (m/s) | 0.052 | 0.050 | 0.046 | 0.077 |
| Mean vertical velocity $\bar{V}_z$ (m/s) | 0.042 | 0.062 | 0.057 | -0.004 |
| Mean horizontal acceleration $\bar{a}_x$ (m/s$^2$) | -0.695 | -0.097 | 0.089 | -1.178 |
| Mean vertical acceleration $\bar{a}_z$ (m/s$^2$) | 1.023 | 0.130 | -0.052 | 4.295 |
| Mean horizontal aerodynamic force $\bar{F}_{ax}$ (μgf) | -0.42 | -0.27 | 0.04 | -0.57 |
| Mean vertical aerodynamic force $\bar{F}_{az}$ (μgf) | 2.71 | 2.38 | 2.98 | 3.26 |
| Horizontal residual error $e_x$ | -10.4% | -9.5% | 0.8% | -12.6% |
| Vertical residual error $e_z$ | 0.9% | -10.5% | 21.5% | -3.3% |
| Mean body mass specific power $\bar{P}$ (W/kg) | 27.6 | 29.5 | 34.0 | 24.4 |
| Peak body mass specific power $P_{max}$ (W/kg) | 109.0 | 115.4 | 164.2 | 90.4 |
| Pitching amplitude $\Delta\chi$ ($^{\circ}$) measured | 29.4 | 24.7 | 28.0 | 32.7 |
| Pitching amplitude $\Delta\chi$ ($^{\circ}$) computed with elytra | 36.2 | 28.2 | 27.8 | 30.2 |
| Pitching amplitude $\Delta\chi$ ($^{\circ}$) computed w/o elytra | 69.8 | 75.3 | 76.6 | 76.0 |

The contribution of the wings, the elytra and the body in the total vertical aerodynamic force is shown in Supplementary Fig. 11. The main contribution is always from the wings. The elytra produce a noticeable positive vertical force towards the end of the morphological downstroke, but this force cancels out with the negative force generated later.

The lift-drag decomposition of the vertical aerodynamic drag shows substantial inter-individual variability, see Supplementary Fig. 12. The instantaneous force is always dominated by the drag, but its large positive and negative contributions cancel each other out after time averaging. Instead, the vertical aerodynamic force due to lift is for most of the time positive. Therefore, the mean vertical aerodynamic force is up to 68%, 84%, 79%, 40% due to the lift and 32%, 16%, 21%, 60% due to the drag for the four beetles, respectively.

**Supplementary Figure 11.**
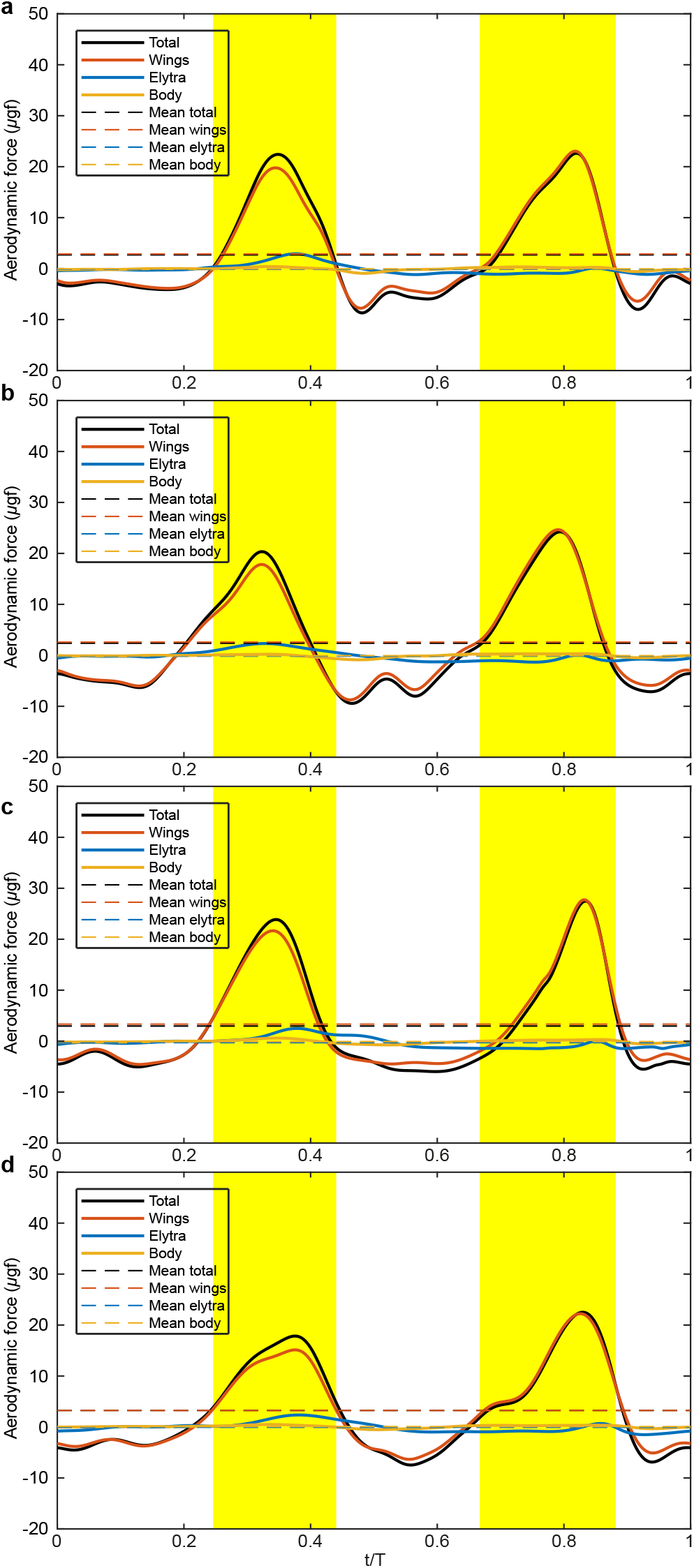
The total vertical aerodynamic force acting on the insect and its breakdown into the vertical component of the aerodynamic force on the pair of wings, the pair of elytra, the body. **a**, PP2. **b**, PP4. **c**, PP5. **d**, PP12.

**Supplementary Figure 12.**
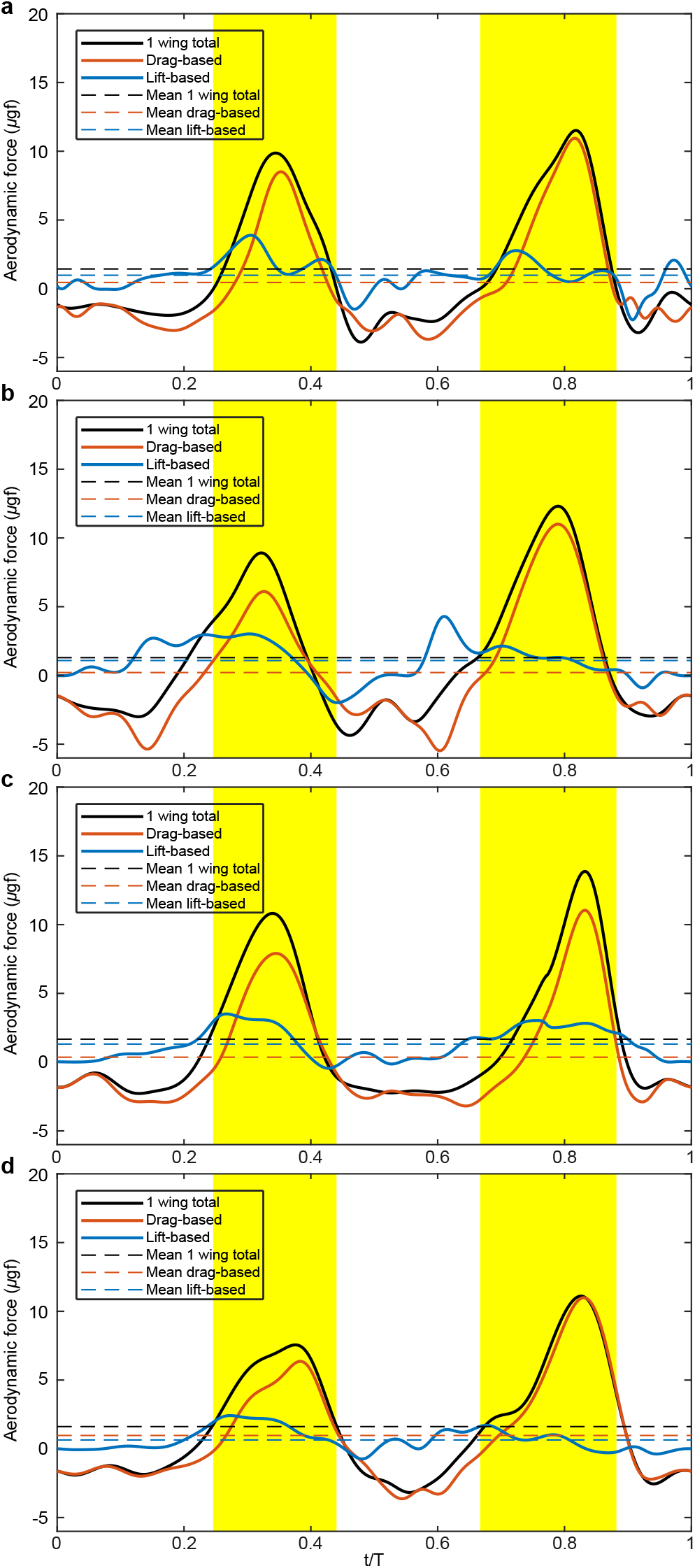
Lift-drag decomposition of the vertical force acting on one wing. **a,** PP2. **b,** PP4. **c,** PP5. **d,** PP12.

The wing tip trajectories shown in Supplementary Fig. 13, coloured according to the sign of the vertical aerodynamic force of the wings, emphasise that the wing beat cycle consists of two power strokes and two recovery strokes. During the power strokes, the wings produce a large aerodynamic force that has an upward component (e.g., at t/T = 0.29 and 0.82). The wing velocity at the same time is downward, because it is tangential to the trajectory and the wing elevation angle decreases during the power stroke. Therefore, large portion of the aerodynamic force can be explained by the drag. However, at t/T = 0.29 for example, the vector diagrams show that the force is not perfectly anti-aligned with the velocity.

It is deflected upwards. This deflection can be explained by the aerodynamic lift which is perpendicular to the direction of wing motion. This positive deflection is present even during the recovery stroke, e.g., at t/T = 0.19, when the aerodynamic force is downward. During the recovery strokes, the wings clap or near-clap and move upwards. This motion generates a downward aerodynamic force. The recovery motion is slow; therefore the aerodynamic force is small in magnitude, but it acts for a longer time. Hence, the net time-average vertical aerodynamic force is an order of magnitude smaller than the peak at the power stroke.

**Supplementary Figure 13.**
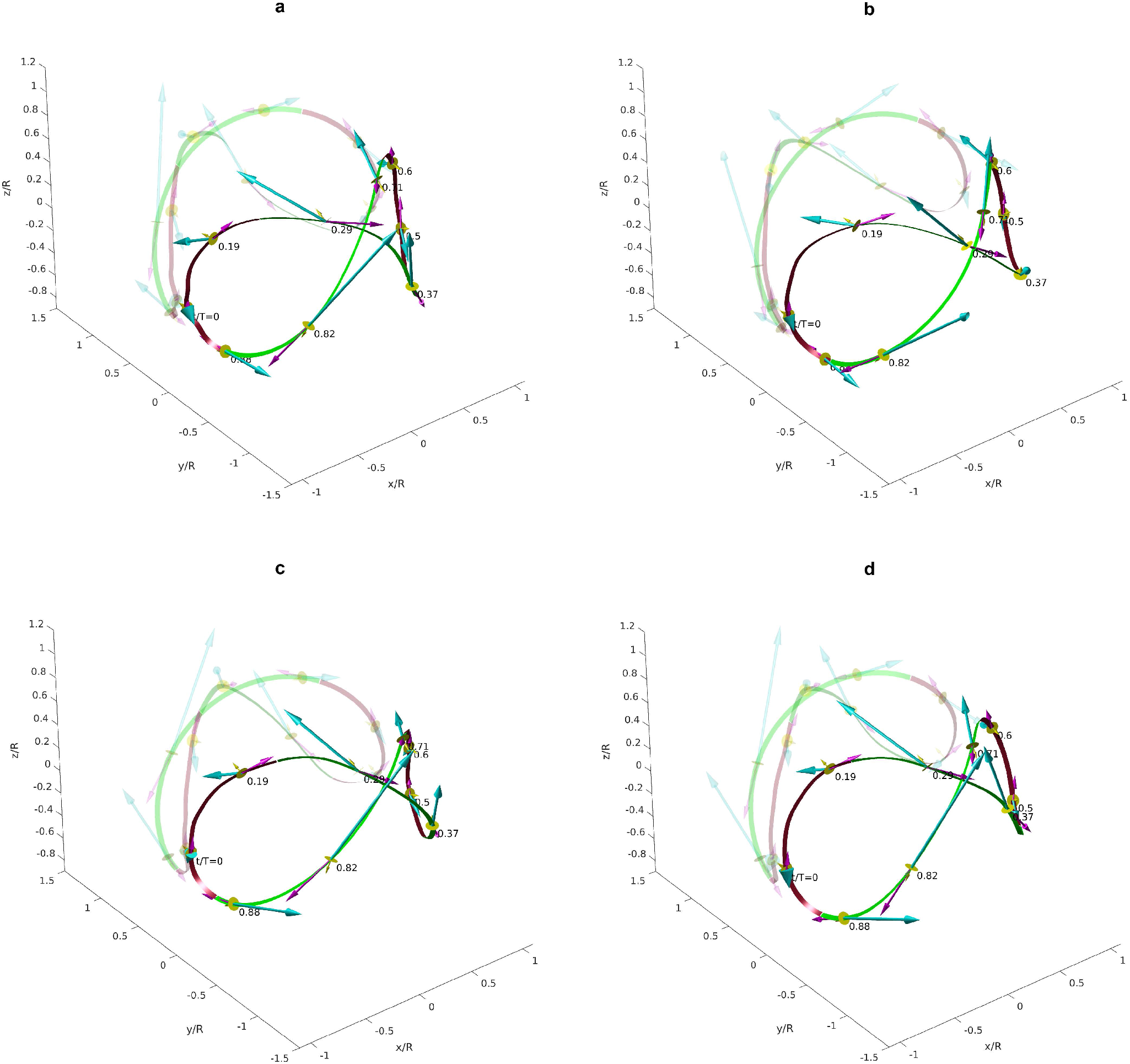
3D reconstruction of the wing-tip trajectories (continuous lines), aerodynamic force vectors (cyan arrows), velocity vectors (magenta arrows) and wing orientation (yellow circles and arrows). Click to view interactive 3D models. **a**, PP2. **b**, PP4. **c**, PP5. **d**, PP12.

The body mass specific mechanical power shown in Supplementary Fig. 14 in all cases peaks during the power strokes. The aerodynamic contribution is dominant. The largest peak is for PP5 (164.2 W/kg) and the smallest peak is for PP12 (90.4 W/kg). The mechanical power essentially remains positive through the entire wing beat cycle period, although occasionally it takes small negative values. For example, its minimum value for PP2 is −5.5 W/kg at t/T = 0.9 and for PP5 it is −8.3 W/kg at t/T = 0.52. Therefore, this type of wing beat motion does not require any substantial elastic energy storage. The mean body mass specific mechanical power is equal to 27.6, 29.5, 34.0 and 24.4 for PP2, PP4, PP5 and PP12, respectively.

The aerodynamic pitching moment that the wings exert on the body (see Supplementary Fig. 15) is consistently negative (nose-up) during the first recovery and power strokes (approximately, morphological downstroke). It becomes positive (nose-down) during the second recovery and power strokes (approximately, morphological upstroke). The inertial pitching moment due to the elytra changes from positive to negative at about t/T = 0.4 when the elytra begin to close to compensate for that changing direction of the aerodynamic pitching moment. The aerodynamic pitching moment of the elytra and the inertial pitching moment of the wings are small.

**Supplementary Figure 14.**
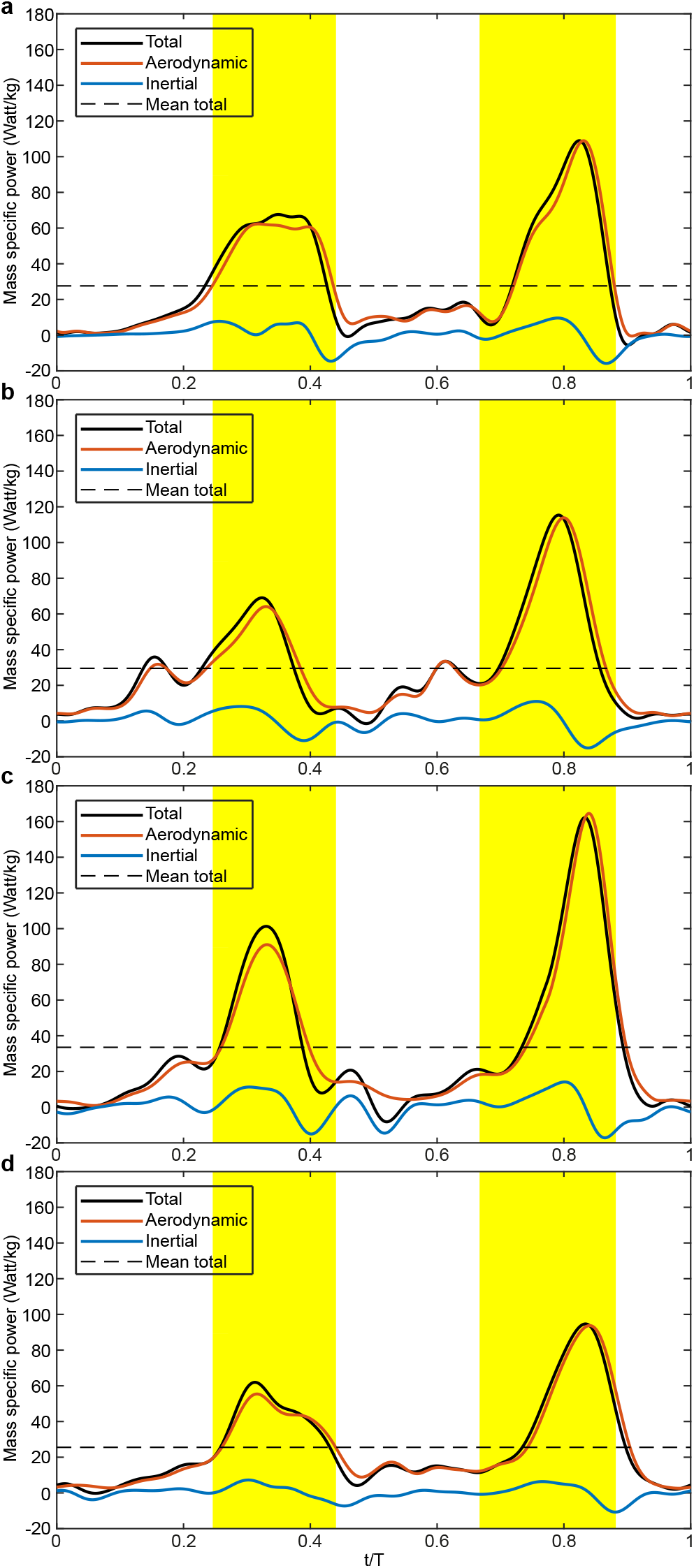
Body mass specific mechanical power components. **a**, PP2. **b**, PP4. **c**, PP5. **d**, PP12.

**Supplementary Figure 15.**
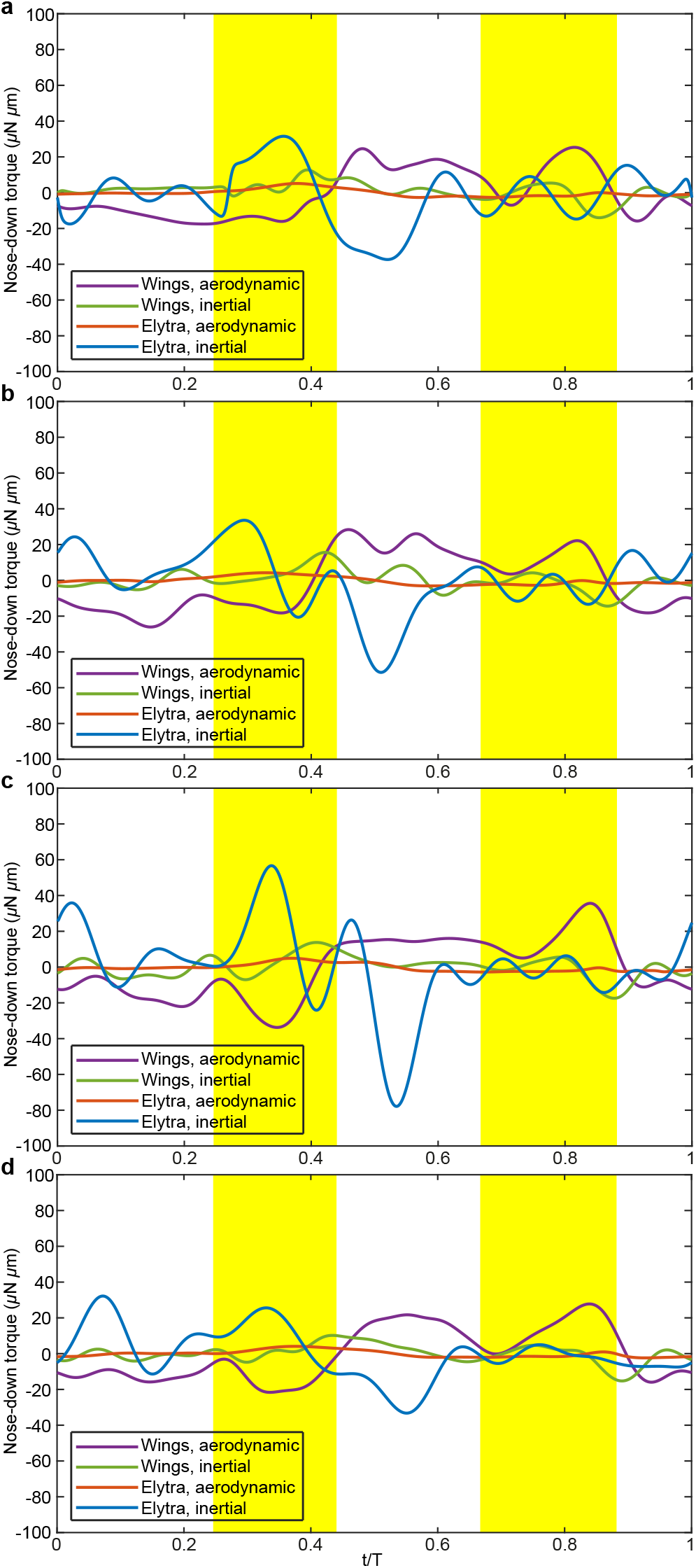
Components of the pitching moment acting on the insect. Positive direction is nose down. **a,** PP2. **b,** PP4. **c,** PP5. **d,** PP12.

### Section 10 – The effect of elytra on body pitch oscillation

To estimate the effect of elytra on the body pitch oscillation, let us consider the angular momentum conservation about the body pitch axis,

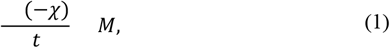

where *M*=*M_ba_*+*M_wi_*+*M_wa_*+*M_ei_*+*M_ea_*, *M_ba_* is the body aerodynamic pitching moment, *M_wi_*, *M_wa_* are the inertial and aerodynamic components of the pitching moment due to the flapping wings and *M_ei_*, *M_ea_* are the inertial and aerodynamic components of the pitching moment due to the elytra, respectively. Note that the “−” sign appears in (1) because the positive direction of *χ* is nose-up (we follow the aeronautical convention) but the positive pitching moment direction is nose-down to be consisted with the right-handed coordinates used in the analysis. Restricting our attention to time-periodic solutions of (1) with the period *T*, we solve it using Fourier transform,

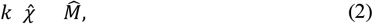

where the hat denotes the Fourier transform of the temporal evolution of a quantity. In the practical calculation, we use the fast Fourier transform in Matlab (MathWorks, U.S.A.). The time evolution *M*(*t*) is sampled on a uniform grid containing *N* = 1024 points, *k* takes integer values from −*N*/2+1 to *N*/2, and we find

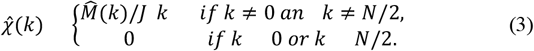

Then, the inverse fast Fourier transform in applied to calculate *χ*(*t*) samples on the same uniform grid of points between 0 and *T*. We mainly want to know the amplitude Δ*χ* = max(*χ*) − min(*χ*).

To avoid additional CFD simulation that would require full fluid-solid coupling, we neglect the dependency of *M_ba_*, *M_wa_* and *M_ia_* on the difference between *χ*(*t*) prescribed in the original CFD simulation and that computed by solving (1). This is justified because the amplitude of the wing motion is much greater than the amplitude of the body motion, and the body and the elytron aerodynamic pitching moments are negligible. The values of Δ*χ* obtained from (1) with *M_ba_*, *M_wi_*, *M_wa_*, *M_ei_* and *M_ea_* taken from the CFD simulations agree well with Δ*χ* measured in the experiment, see Supplementary Table 3. This additionally validates the CFD model.

Supplementary Figure 15 further demonstrates that *M_ea_* is negligible. But the inertial pitching moment due to the elytra *M_ei_* is essential. If we fully neglect it, substituting *M* = *M_ba_* + *M_wi_* + *M_wa_* into (1), we find values of Δ*χ* that are 1.9 to 2.7 times larger than in the previous cases with the elytra. The maximum body pitching angle max(*χ*) becomes greater than 90°. We therefore conclude that the elytra act as inertial brakes that prevent overturning.

### Section 11 – Numerical convergence

Additional numerical simulations have been carried out to evaluate the accuracy of the CFD simulations. In addition to the original simulation of PP2 with the maximum number of refinement levels *J_max_* = 9, we have repeated similar computations with *J_max_* = 8 and *J_max_* = 7. These new runs have a coarser grid resolution in the neighbourhood of the solid boundaries (see the values of Δ*x_min_*/*R* in Supplementary Table 3).

Supplementary Figure 16 shows the time variation of the aerodynamic force components and the body mass specific aerodynamic power in these three computations. All three show similar qualitative trends. The results for *J_max_* = 8 and *J_ma x_*= 9 are close enough for making quantitative conclusions. Thus, the difference between the mean vertical force computed with *J_max_* = 8 and with *J_max_* = 9 is less than 9% of the body weight. The horizontal force difference is less than 1% of the body weight. The mean aerodynamic power differs by less than 4%.

**Supplementary Figure 16.**
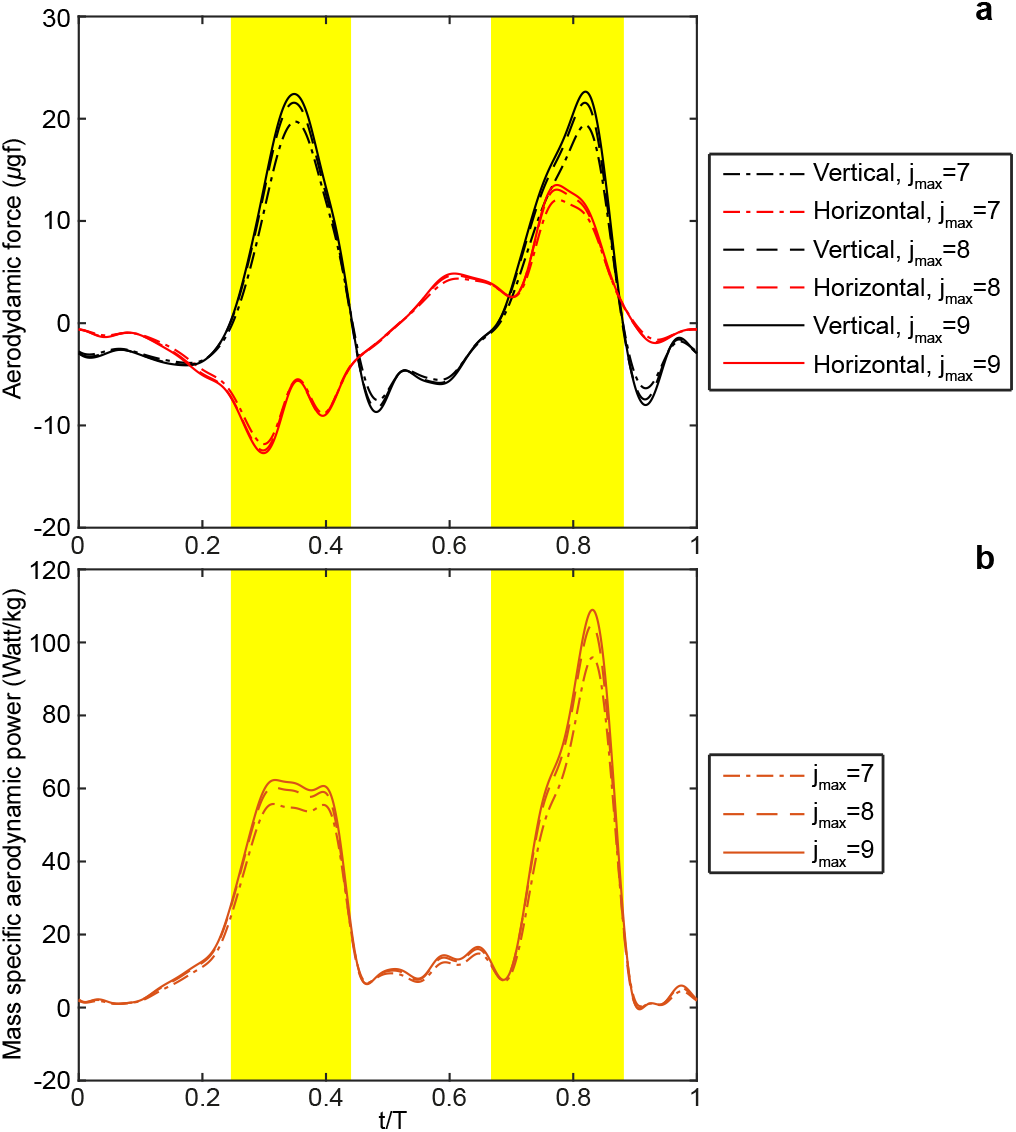
Aerodynamic performance of PP2 obtained from three different simulations using three different values of the maximum refinement level. **a,** Aerodynamic force. **b,** Mass specific aerodynamic power.

### Section 12 – On the fluid-dynamic added mass effect

It is common to use the concept of added mass to describe the acceleration reaction of a solid body moving in a fluid. An order-of-magnitude estimate can be obtained by the following formula^8^:

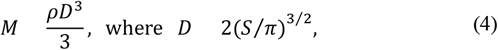

*M* is the added mass; *ρ* is the air density (1.197 kg/m^3^); *S* is the wing area. Assuming the added air mass of the bristled wing (for its low leakiness) is about the same as the added air mass of the membranous wing model, we find *M*=0.0233 μg, which is close to the mass of the bristled wing. We do not develop further on this question, because the acceleration reaction at *Re* ≈ 10 strongly depends on the flow velocity, which undermines the practicality of the added mass concept.

**Supplementary Table 3.**
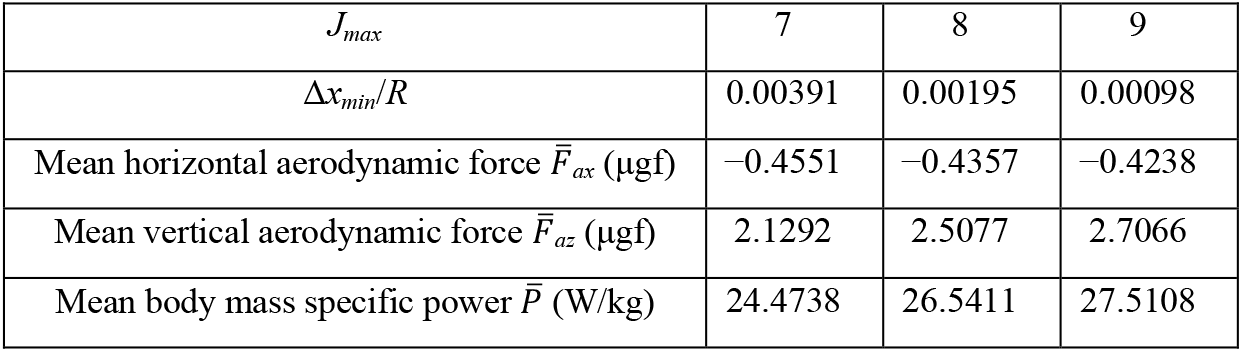
Numerical convergence data.

